# Occurrence of non-western magic in the European brain

**DOI:** 10.1101/395285

**Authors:** Jan Willem Koten, André Schüppen, Vinod Kumar, Guilherme Wood

## Abstract

Timecourses that exhibit identical behaviour at distinct measurement occasions are reliable. Voodoo connectivity occurs when connectivity among brain regions exceeds within subject timecourse reliability. Thus, timecourse reliability limits the true detectable connectivity. We reproduced a working memory related connectome consisting of 561 paths obtained from 67 individuals. We tested >100000 fc-MRI pipelines and show that Savitzky Golay (SG) filters maximize true connectivity while conserving cognitively relevant changes of signals. This is noteworthy for approaches that focus on rapidly changing aspects of connectomes. Furthermore, SG filters detect zombie activity. These “resting state oscillations” are not under human control and contaminate working state signals. SPM pipelines exhibit more voodoo connectivity than SG pipelines. With the SPM pipeline, we observed a connectivity of r=0.44 and a poor true connectivity of r=0.23, but with the SG pipeline we observed a connectivity of r=0.59 and a fair true connectivity of r=0.43. The number of paths detected with fair true connectivity (r >0.4) was 4 for the SPM pipeline but 352 for the SG based pipeline. However, superior statistical properties of SG pipelines may not reflect neural reality. Hence, causal external validation of fc-MRI pipelines is crucial. Without such studies, different pipelines produce at best “alternative maps”.

## ERRATTA

The first author of this text is a slightly outdated individual without a portable telephone. I stem from a time when people enjoyed colourful language. However, I feel somewhat uncomfortable with the fact that small fragments of the abstract are pushed through the internet with breath taking speed without the context. I am not sure if people understand my paper correctly. I painfully realize that a modern paper should be twitter compatible, which the first version of the paper was obviously not. I would like to excuse myself for my shortcomings. The expression ‘SPM pipeline’ in the abstract must be understood within the context of the method section of the paper. Maybe it sounds a bit old fashioned and silly but I would appreciate it if people would read the method section. I could be wrong after all. I thank all the twitter people for their interest. Maybe it is just an enthusiastic comeback of the telegram. Albeit, a bit fast for my head.

To avoid any confusion I will give a new abstract that is hopefully twitter compatible.

Sincerely,

Jan Willem Koten

Timecourses that exhibit identical behaviour at distinct measurement occasions are reliable. Voodoo connectivity occurs when connectivity among brain regions exceeds within subject timecourse reliability. Thus, timecourse reliability limits the true detectable connectivity. We reproduced a working memory related connectome consisting of 561 paths obtained from 67 individuals. We tested >100000 fc-MRI pipelines and show that Savitzky Golay (SG) filters maximize true connectivity while conserving cognitively relevant changes of signals. This is noteworthy for approaches that focus on rapidly changing aspects of connectomes. Furthermore, SG filters detect zombie activity. These resting state oscillations are not under human control and contaminate working state signals. SPM filters exhibit more voodoo connectivity than SG filters. With the SPM filter based pipeline, we observed a connectivity of r=0.44 and a poor true connectivity of r=0.23, but with the SG pipeline we observed a connectivity of r=0.59 and a fair true connectivity of r=0.43. The number of paths detected with fair true connectivity (r >0.4) was 4 for the pipeline that was based on the SPM filter but 352 for the SG pipeline. However, superior statistical properties of SG pipelines may not reflect neural reality. Hence, causal external validation of fc-MRI pipelines is crucial. Without such studies, different pipelines produce at best “alternative maps”.

## BACKGROUND

One of the most read papers in science is entitled “*Why most published research findings are false”* (1). Empirical investigations indeed show that science is plagued by a reproducibility crisis that may cost US$28.000.000.000 in the US alone (2-4). The reproducibility crisis is partly caused by measurement error, misuse of significance tests, poor research methods and by “*senior people who impose perverse incentives on scientists* “(5-14). Scientists such as Cohen, Meehl and Carver have argued that statistical significance is not equivalent with scientific significance (6-8). Cohen was concerned particularly with the problems that may arise from statistical hypothesis testing and quoted Meehl in his seminal paper entitled the “*The Earth Is Round (P-Less-Than.05)*”. Meehl described significance testing as: “*a potent but sterile intellectual rake who leaves in his merry path a long train of ravished maidens but no viable scientific offspring*”(6). The low level of *“scientific offspring”* is partly related to the misconception that statistical significance is synonymous with reproducibility. Carver described this “*replicability or reliability fantasy*” in his classic paper “*the case against significance testing*”(8). Recently, Colquhoun argued that the terms *‘significant’ and ‘non-significant’ should never be used* (9). This is reasonable given that most reported studies in the medical field are statistically significant while reproducibility rates are low (10). Cohen, Carver and other researchers have argued that reproducibility, point estimation and confidence intervals are at the core of science (6-9).

Currently, functional connectivity MRI studies follow the scientific main stream. The emphasis is on significance testing while ***true*** point estimates of connectivity strength are remarkably absent from the literature. In this study we try to obtain true estimates of connectivity strength. This is challenging because true connectivity strength depends on timecourse reliability that in turn depends on signal processing methods. Thus, true connectivity strength, timecourse reliability and signal processing methods are mutually depended phenomena that are hard to separate from each other. So far, numerous studies assessed the reliability of brain activity and connectivity on group- and single subject level with a rich arsenal of methods (15-30). Recent research shows that reliability of resting connectivity metrics is sensitive to preprocessing techniques (15, 16, 26). But established methods are not suitable to study the effects of signal processing on true connectivity. True connectivity is closely related to the voodoo correlation phenomenon that was described in the paper “*Voodoo Correlations in Social Neuroscience”.* The title of this paper led to considerable uproar in the community and the authors were politely requested to rephrase their paper in politically acceptable terms (31). Currently the phrase voodoo correlation has established itself as scientific nomenclature despite attempts to censor it (32-35). To refresh the mind of the reader we present the original statistic arguments of Nunnally.

***“It is a statistical fact (first noted by researchers in the field of classical psychometric test theory) that the strength of the correlation observed between Measures A and B (rObserved*_*A*_,*Observed*_*B*_*) reflects not only the strength of the relationship between the traits underlying A and B (r*_*A,B*_*), but also the reliability of the measures of A and B (reliability* _*A*_ *and reliability* _*B*_, *respectively).***

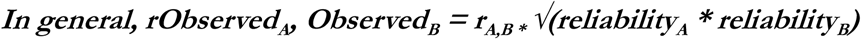

***Thus, the reliabilities of two measures provide an upper bound on the possible correlation that can be observed between the two measures. This is the case because the correlation coefficient is defined as the ratio between the covariance of two measures and the product of their standard deviations: r*_*x,y*_ *= σ*_*xy*_ */σ*_*x\**_*σ*_*y.*_ *Real-world measurements will be corrupted by (independent) noise, thus the standard deviations of the measured distributions will be increased by the additional noise (with a magnitude assessed by the measure’s reliability). This will make the measured correlation lower than the true underlying correlation by a factor equal to the geometric mean of reliabilities. Thus, the reliabilities of two measures provide an upper bound on the possible correlation that can be observed between the two measures “*(36)**

The original “voodoo correlation” paper criticized fMRI group studies that correlate the bold response with external measures of behaviour. But the very same critique holds true for functional connectivity studies at the single subject level. I.E. the height of a functional connectivity correlation between two regions A and B cannot exceed the test retest reliability correlation of underlying timecourses. It is inherently impossible to estimate test retest reliability of resting state timecourses because the latter are not synchronized by an external stimulus. Consequently, we will not further discuss resting state studies since they present incommensurable problems regarding signal reliability.

Currently, more than 40.000 fMRI paper have been reported but studies that report reliability of task driven timecourses on the within subject level remain very rare (11, 22). It is likely that functional connectivity at the single path single subject level is voodoo given that within subject test retest reliability of timecourses is slightly disappointing. Mean voxel wise timecourse reliability estimated from the full brain is around r = 0.1 (22). At best correlations of 0.25 have been reported in motor areas that were activated during a motor task (22). This suggests that true connectivity among region A and B cannot exceed 0.25. The poor timecourse reliability is most likely related to the poor signal to noise ratios of MRI scanners (37). The latter obscures timecourse reliability. Over the years, several preprocessing pipelines have been developed to reduce noise. In classic versions of the SPM package discrete cosine transforms were used to remove slow trends form the signal (128seconds) while HRF or Gaussian smoothing kernels (around 2.4sec) were used to remove high frequency noise (38). But in particularly low pass filters may boost auto correlations of timecourses and therefore destroy the temporal resolution of a timecourse (39-41). This is in particular problematic for event related designs that ideally rely on timecourses that exhibit higher temporal resolution. Consequently, low pass filters have been removed from current versions of SPM.

The following question arises: Is it possible to develop low pass filters that do justice to the “natural” autocorrelation structure of a timecourse, and if yes, what is the true auto correlation of an fMRI timecourse? Originally, the so called HRF function was derived by averaging the BOLD responses of events induced experimentally. This event related average is a denoised approximation of what the ideal BOLD response for a given brain region and individual might look like. Empirical predictors created from slow event related designs are obtained by concatenating event related averages in agreement with the number of times that a psychological event took place. Empirical predictors might reflect the true auto correlation estimate of a timecourse.

It has been shown that Savitzky-Golay (SG) filters conserve much of the original signal contour(42). Hence SG filters may preserve much of the true auto correlation structures of fMRI signals (42). Although SG filters are used in many branches of science - such as chemistry, medicine, ecology etc. - they have rarely been used in the context of fMRI. One fMRI study used an SG filter to estimate the contrast to noise ratio of a timecourse (43). The second study used SG filters to reconstruct changes in the phase of MRI signals obtained at 7T. These “phase timecourses” can be used to suppress spurious BOLD activations stemming from vessels and draining veins while conserving signal changes from microvascular effects (44).

The SG algorithm consists of three components: (i) a sliding window of size *m* is selected; (ii) a polynomial function is fitted to the observed data within the window. Finally, (iii) the data point observed experimentally at the centre of the window is replaced by the predicted polynomial value. Next, the window is moved by one sampling interval. Subsequently, procedures i-iii, are applied to the observed data of the shifted window. The number of time points of the window *m* must be uneven because it should have a clearly defined centre i.e. the number of points at the left and right side of the centre should be identical. It is obvious that the order of the fitted polynomial can never exceed the number of degrees of freedom, which are given by the size of the window -1. Thus, the highest possible polynomial order is *m-1*. It is also clear that sliding windows cause problems at the beginning and the end of timecourses. An accepted solution is to artificially extended data by adding, in reverse order, copies of the first (*m* - 1)/2 points at the beginning and copies of the last (*m* - 1)/2 points at the end (42). We have followed this procedure it the present study. The main problem of the SG filter is to find the optimal combination of window size and polynomial order. In principle, two methods are available: brute force search in empirical data or computer simulations in synthetic data. Computer simulations are usually based on a synthetic signal that is contaminated with synthetic noise. This method is powerful because it specifically tests if a filter can identify the “known” synthetic noise correctly. The elegant computer simulation method assumes that synthetic noise is a realistic reflection of the noise as it occurs in real fMRI data. But the latter is most likely not the case because distinct sources of noise exhibit complex behaviour that may also vary between subjects and brain regions(45). For this reason, it has been suggested that optimal SG filters may be obtained through brute force methods in empirical data (44).

Here, we investigate if SG filters minimize voodoo connectivity and spurious auto correlations. In a first step we determined optimal SG parameters in a verbal working memory task through brute force methods. We assume that a good detrending and low-pass filtering system should maximize the correlation between an observed timecourse and a predicted timecourse. In this study, the predicted timecourse was estimated with the same psychological experiment measured at a different session. This avoids problems of circularity. In second step we validated our SG pipeline in a spatial working memory task and compared the result with classic fMRI pipelines. The stimuli of the cognitive tasks that were used for SG filter optimization differed from those of the validation study to avoid circularity. Finally, we investigated how task driven connectomes react when resting baseline conditions are removed from timecourses.

## METHODS

### Scanner parameters

MRI scans were performed on a 3T Siemens Magnetom Skyra (Siemens Medical Systems, Erlangen, Germany) equipped with a 32-channel head coil. We used a 3D-MPRAGE sequence (176 slices per slab, FOV = 256 mm, TR = 2530 ms, TE = 2.07 ms, TI = 900 ms, Flip angle = 9°, voxel size = 1mm isotropic). Functional imaging data were obtained using a Siemens Grappa parallel acquisition scheme with pat factor 2; using following parameters Flip Angle 72 degrees, TR = 1240 ms, TE = 30 ms. Volume dimensions were 64*64*23, with voxel resolution 4*4*4 mm with a gap of 10% For the working state analysis in total 487 volumes were obtained per task per run.

### Participants, working memory task, Coregistration and Timecourse extraction

The 67 individuals studied, varied in age and academic attainment and are representative of the Austrian population. Participants performed two tasks: A verbal working memory task that incorporated a numerical distractor task and a spatial working memory task that incorporated a numerical distractor task. We used a slightly larger jitter as usually advised. The main reason is to demonstrate that SG filters can cope with highly jittered timecourses. Details of the task are found in the Supporting Material (SM) and Figure 1. We obtained test and retest data from both tasks. Test and retest sessions were interrupted by a pause of at least one hour outside the scanner. Nuisance timecourses were obtained using standard FreeSurfer options (https://surfer.nmr.mgh.harvard.edu/). Grey matter timecourses were brought into FS_average space using spherical alignment methods. Subsequently, 34 working memory related regions of interest stemming from a meta-analysis were projected onto FS_average mesh space (46). Timecourses were extracted from 34 patches of interest on the surface and subjected to further analysis. Next, we visualized the resulting connectomes with connectome viewer software package (http://nica.net.cn/achievements/t20120530_1132.htm). We used the coordinates of meta-analysis for this purpose. It is important to state that for visualisation purposes connectomes where shown in 3d space. However data were in reality in 2d spherical aligned space. Further details are given in the Supporting materials.

**Figure 1:**
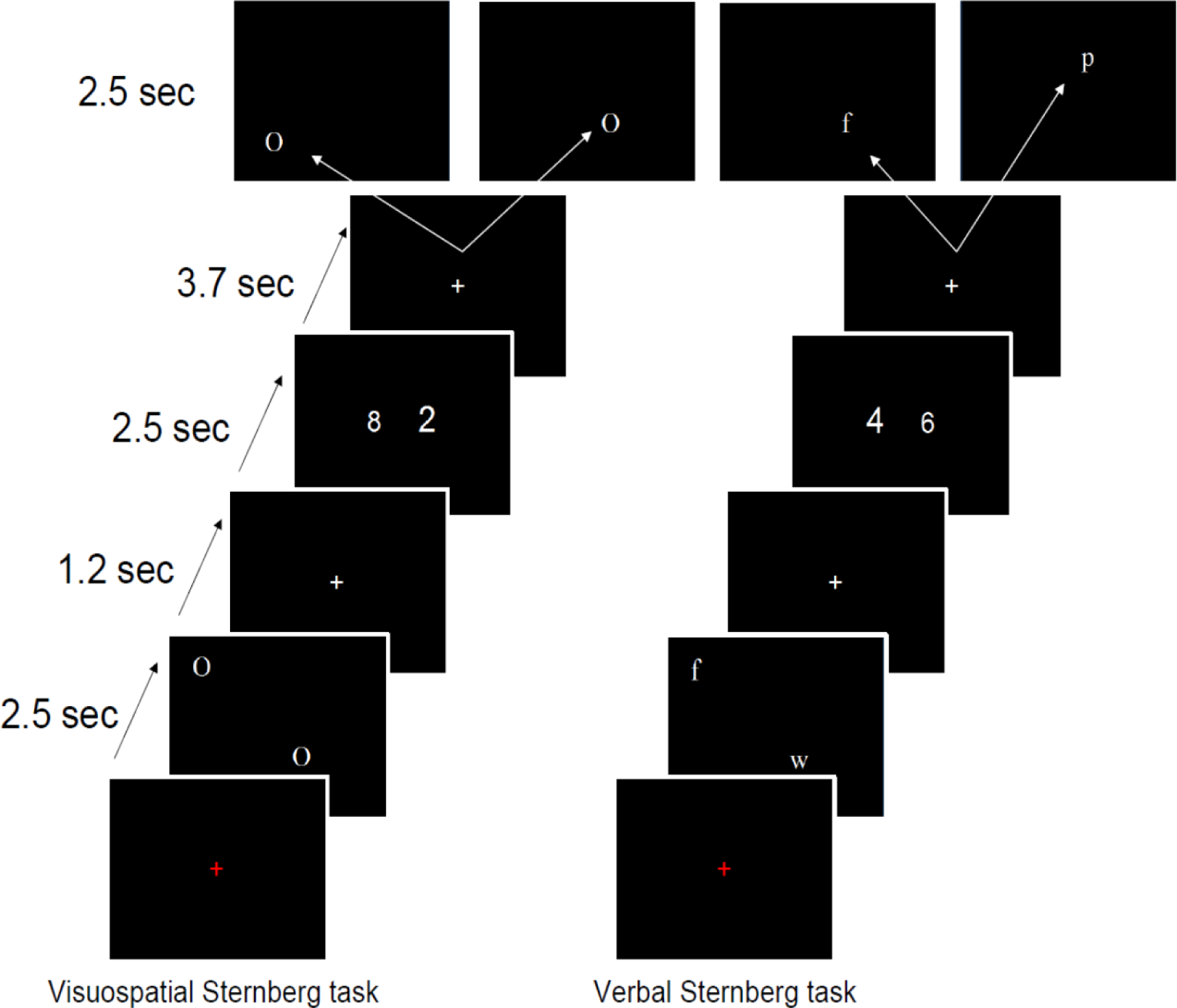
This figure illustrates details of the spatial and verbal working memory tasks that were used for the optimization and validation experiment respectively. The exact duration of the stimulus is depicted on the left side. Mark that the resting baseline depicted with a red fixation cross was jittered. It varied roughly between 11.16 sec. and 14.88 sec. Left side, individuals had to memorize the spatial position of two “O’s” subsequently they had to identify the larger numerosity of two Arabic numbers that were different in physical size and press a button with the left or right index finger according to the side were the larger numerosity was found. Finally participants had to verify whether the spatial position of a newly presented “O” was in agreement with the previously memorized “O” set. If this was the case participants pressed the right button if not the left. Right side, the verbal memory task followed the same regime. But in this case participants had to verify the identity of the letter but not the spatial position of the letter.

### General overview of the experiment and MATLAB^®^ packages

In this experiment, SG filtering was applied to remove slow signal drifts (detrending), and high frequency noise from fMRI data. The method was developed on the basis of in house MATLAB^®^ scripts - that comprise 1000 lines of novel code - and several toolboxes of the MATLAB^®^ family. The aim of the distinct scripts was to investigate the different aspects of the filter. The first script “find_filter.m” was used to optimize filter parameters for detrending and low pass filters; we will refer to this as optimization experiment. The script was applied to a data set containing a verbal working memory experiment that consisted of a test and a retest phase. The second script “test_filter.m” was used to evaluate the effects of signal processing methods on test retest reliability, connectivity, true connectivity and auto correlation of timecourses. Again, this spatial working memory experiment consisted of a test and a retest phase. We will refer to this as validation experiment. The rationale behind this design was the following. Optimization and validation of the filter parameters need to be executed with two distinct experiments to avoid circularity (verbal and spatial working memory). However, the temporal structure at which the psychological events are presented need to be similar for the two distinct experiments. Otherwise, the filter parameters obtained during the optimization experiment might not work for the validation experiment.

### Methods for Optimization experiment

Prior to the optimization procedures we performed slice scan time correction and motion correction. Subsequently, we constructed predictor timecourses, which were based on event related averages from a test run. Next, we correlated the SG filtered timecourse obtained from a retest run with the predictor timecourse. We performed a correlation procedure for the whole window and polynomial space of the SG filter and determined the iteration at which maximal correlations between predicted and observed timecourses was obtained. Usually, the correlation between phenomena A and phenomena B is identical with its inverse. However, in this study the event related average of the test run is not identical with the event related average of the retest run. While the same logic holds true for the observed (filtered) data of the respective runs. Hence, the predictor timecourse was estimated from test run and correlated with retest run and vice versa. This resulted in 2 runs*34 nodes*67participants=4556 correlation for every iteration step. Subsequently, we averaged the correlations observed per iteration. The filter parameters that maximized the correlation between observed and predicted timecourse were taken to be the optimal parameters which were tested for their effects in a distinct cognitive experiment. We used SG filters to remove slow trends and high frequency noise. For this reason, an incremental strategy was used. In a first phase we developed the optimal SG filter parameters for detrending purposes (high pass filter). In a second phase, the optimal detrending parameters were used to create the optimal low pass filter parameters.

Phase 1: The predicted timecourses were obtained from denoised data, while observed timecourses were obtained from denoised and SG detrended data. The pipeline is depicted in Figure 2. The predicted timecourse was constructed by averaging 24 task related timecourse sections. The resulting event related average, which is the idealized hemodynamic response for a given node in a given person, was repeated 24 times. As described in the previous method section, this experiment was jittered. Hence, the length of resting pauses between working memory tasks were irregular. Consequently, we extracted 24 events of identical length from the test and retest signal and omitted parts of the timecourse that were not shared. In Figure 3, we show how the event related average was constructed. For the detrending optimization procedure, the whole window space from 3 to 487 TR’s was investigated in steps of 2. For each window size, the whole polynomial order space from one to window size -1 was investigated. Finally, we estimated the autocorrelations of the predictor timecourses that were denoised and detrended with optimal SG filters within a GLM framework. Autocorrelations included lags 1 to 4.

**Figure 2:**
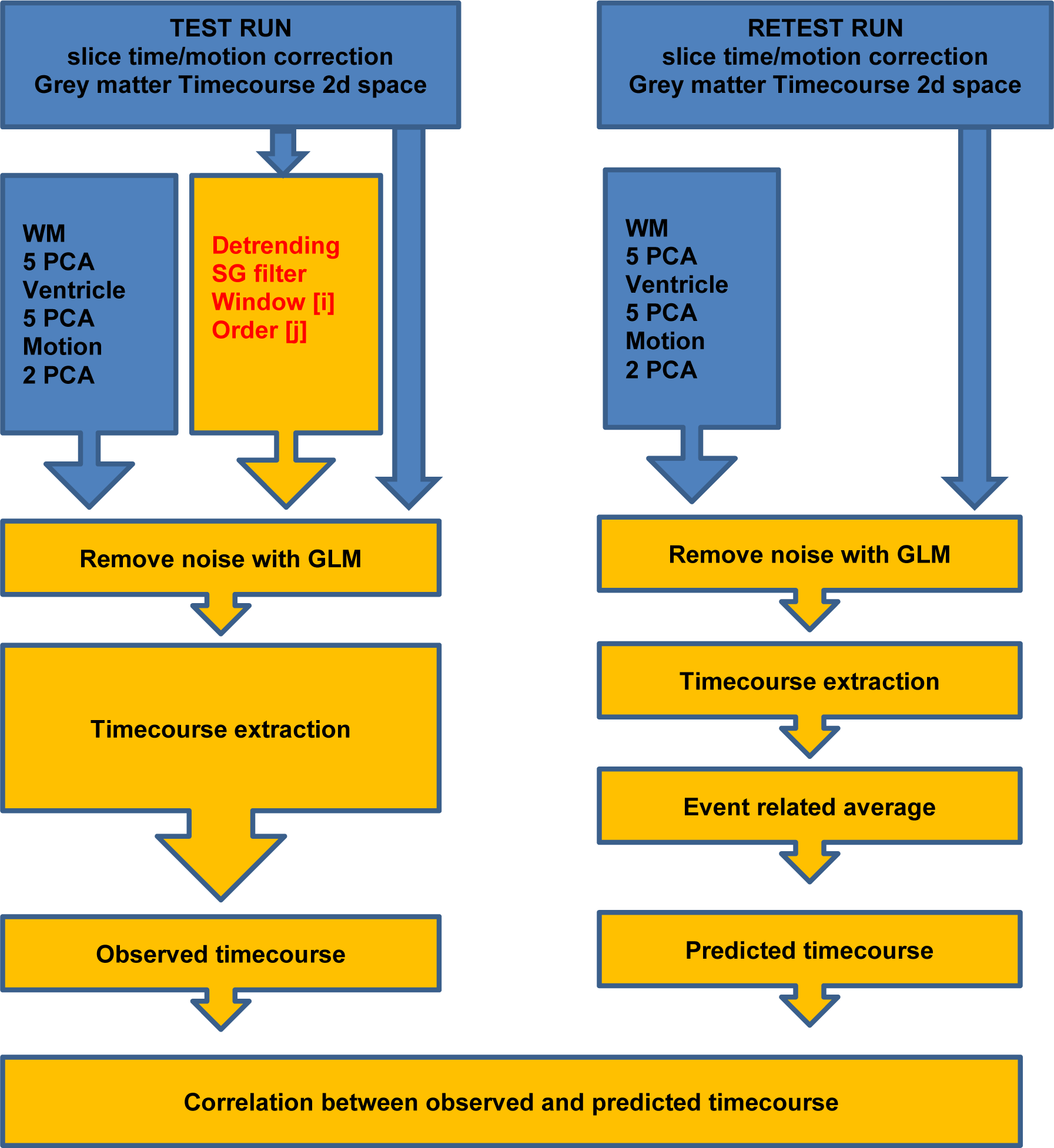
This figure shows the pipeline for the first phase of the optimization experiment that aimed at finding the optimal SG filter parameters for detrending. Red text refers to SG parameters that were changed at every iteration step. This means that the whole window space from 3 to 487 TR’s was investigated in steps of 2. For each window size [i] the whole polynomial order space [j] from one to window size -1 was investigated. Blue parts of the graph refer to operations that were executed using the FreeSurfer package while the yellow parts refer to in house MATLAB routines.

**Figure 3:**
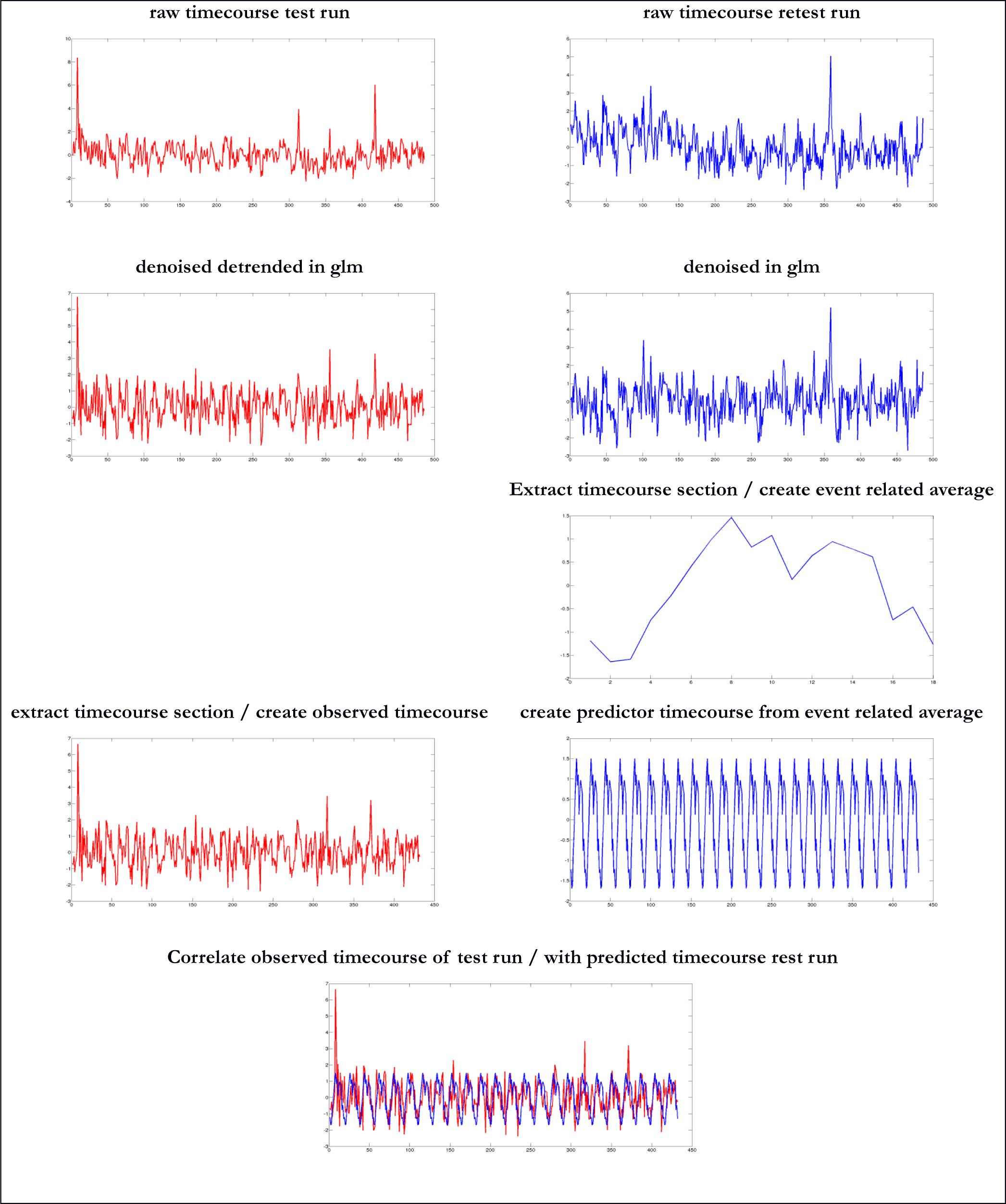
This figure shows the effects of preprocessing on the timecourse as executed in the first phase of the optimization experiment depicted in Figure2 that aimed at finding optimal detrending parameters.

Phase 2, the predicted timecourse were obtained from denoised and optimal SG detrended data, while observed timecourses were obtained from denoised, SG detrended data and SG low pass filtered data. For this purpose we selected a quasi-optimal SG filter with a window of 69 TRs and a polynomial order of 6 (69/6) and an optimal SG filter with a window of 311 TRs and a polynomial order of 40 (311/40). The rationale behind this operation is to show that suboptimal filters that are suitable for timecourses with a small number of observations may deliver results that are comparable to optimal filters that require a large number of observations. The exact pipeline and its effect on the timecourse are shown in Figure 4 and Figure 5. In short, we investigated the whole window space from 3 to 487 TR’s in steps of 2. For each window size the whole polynomial order space from one to window size -1 was investigated. However, we did not investigate polynomial orders larger than 50 when using the low pass filters. Again, correlations between predicted and observed timecourses were estimated. In addition, autocorrelations of observed timecourses sections were estimated in every single iteration step. These included the lag 1-4 autocorrelations.

**Figure 4:**
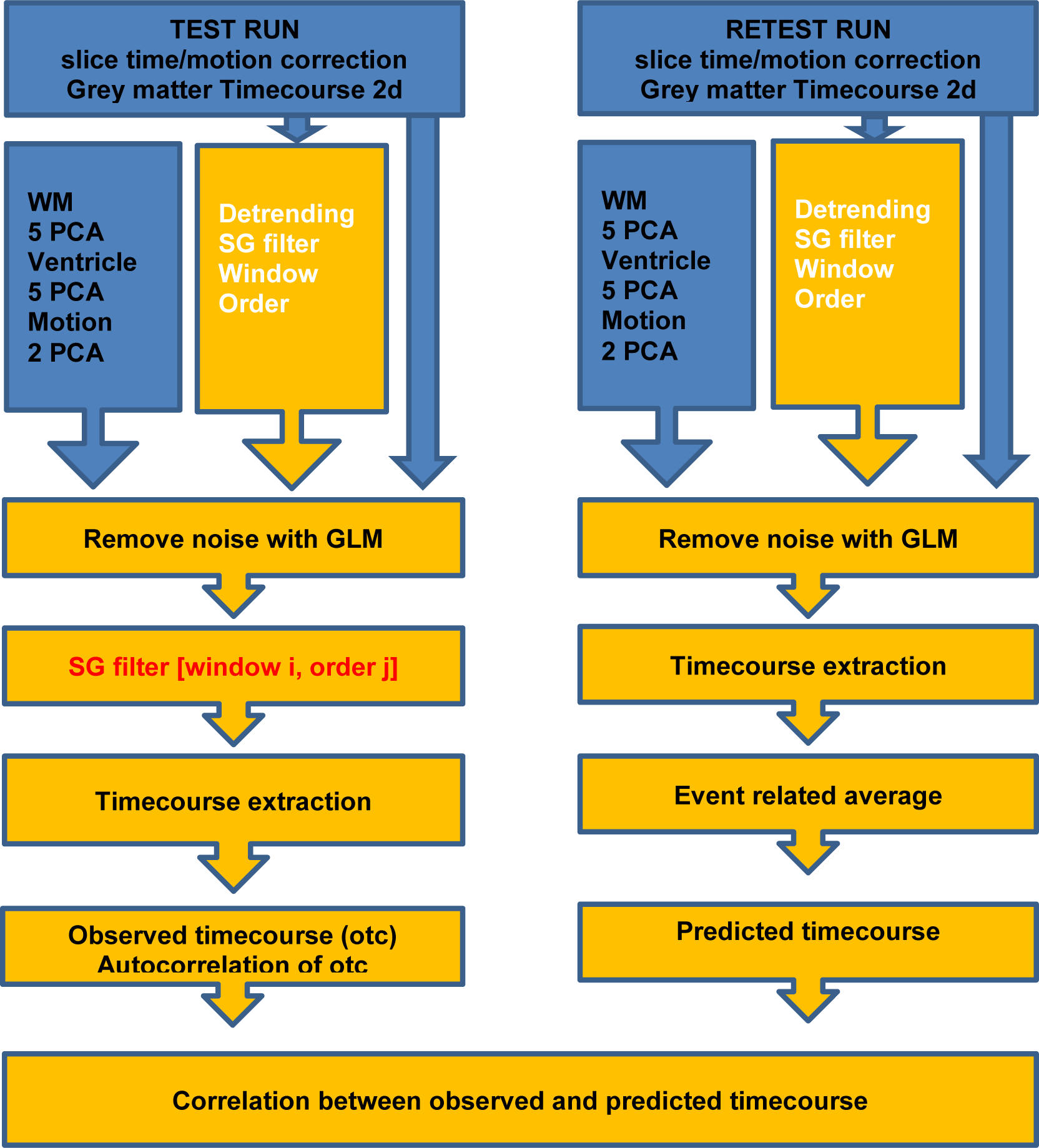
This figure shows the pipeline for the second phase of the optimization experiment that aimed at finding the optimal SG parameters for low pass filtering. Red text refers to SG parameters that were changed at every iteration step. This means that the whole window space from 3 to 487 TR’s was investigated in steps of 2. For each window size [i] the whole polynomial order space [j] from one to window size -1 was investigated. However polynomial orders larger 50 were not investigated. Blue parts of the graph refer to operations that were executed using the FreeSurfer package while the yellow parts refer to in house MATLAB routines.

**Figure 5:**
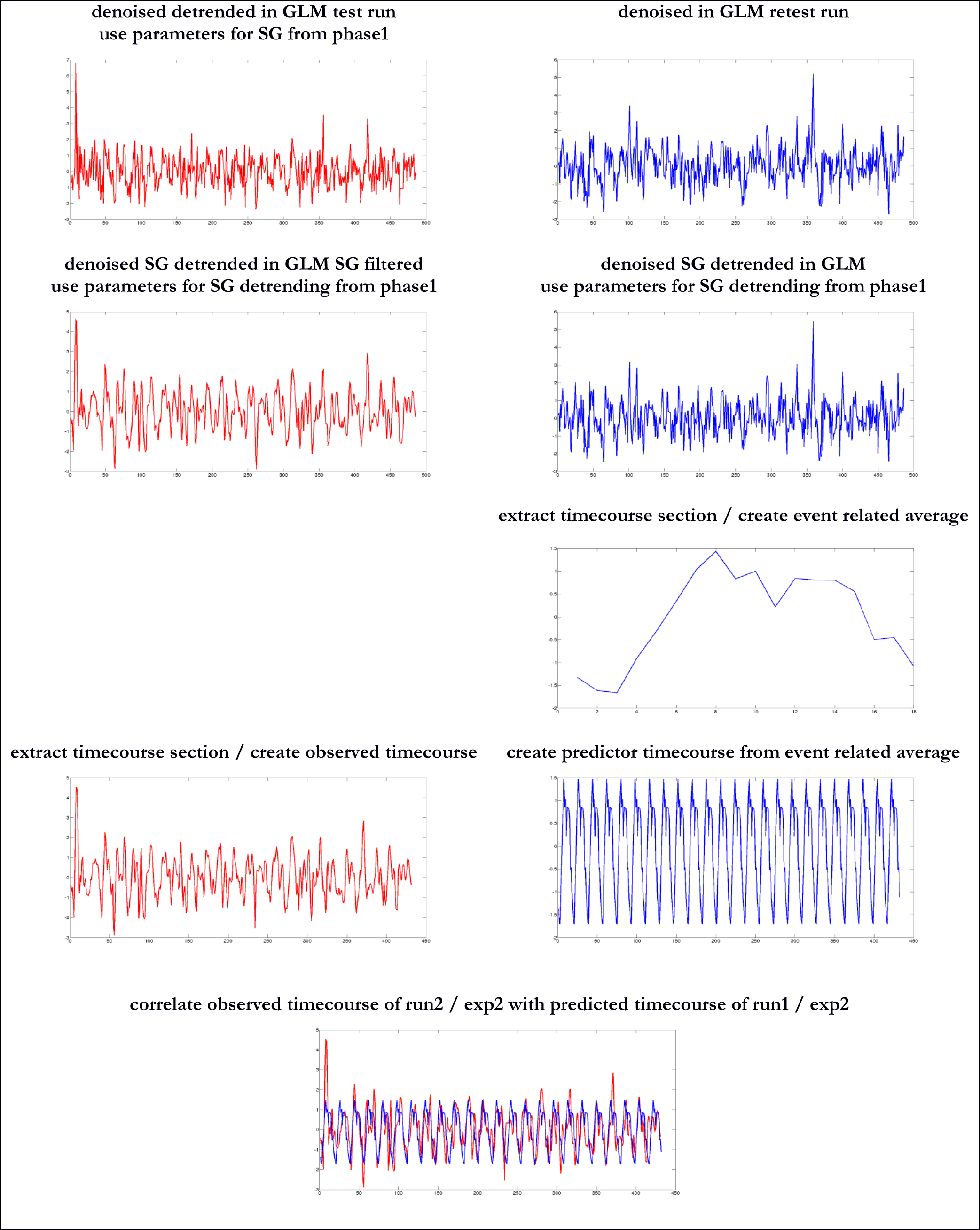
This figure shows the effects of preprocessing on the timecourse as executed in the second phase of the optimization experiment depicted in Figure 4 that aimed at finding the optimal low pass filter.

### Root mean square error of predicted and observed auto correlations

As mentioned in the introduction we assume that predicted timecourses that are based on event related averages - constructed from timecourses that were denoised and detrended within a GLM frame work - represent the true auto correlational structure of a timecourse. We used a SG (311/40) detrending filter for this purpose. Subsequently, the autocorrelations estimated from 34 nodes*67 participants were averaged per session and used for root mean square error (RMSE) estimation. We estimated the auto correlations of observed timecourse sections at every single iteration step during the second phase of the optimization experiment. Subsequently, the autocorrelations of 34 nodes*67 participants were averaged per iteration. In a next step, we estimated RMSE per iteration from the 4 autocorrelations of the predicted timecourses and 4 autocorrelations of observed timecourses. Finally, the correlations between the predicted and observed timecourses were masked with RMSE<0.1. We assume that the resulting graph reflects filters that maximize the correlation between predicted and observed timecourses while autocorrelations of timecourses are kept at an acceptable level.

### Methods for validation experiment

Contrary to the default FreeSurfer FsFast pipeline, we performed slice scan time correction prior to the motion correction. The motion parameters that were created during motion correction underwent a principal component analysis. The first two principal components of the motion parameters were extracted from FsFast/mcprextreg file and used as nuisance regressors. The “eroded” white matter and ventricle timecourses underwent principal component analysis. We used the FsFast default settings and extracted the top five components of the white matter and ventricles respectively. Next, either a quasi-optimal SG filter with a window of 69 TRs and a polynomial order of 6 (69/6) or an optimal SG filter with a window of 311 TRs and a polynomial order of 40 (311/40) or a SPM high pass filter (128 secs. > 0.0078 Hz) was run over the timecourses. The resulting timecourses were used to remove slow trends from the timecourse (detrending). The 2 motion components together with the 10 subcortical nuisance timecourses as well as the slow trend of the grey matter timecourse were imported into MATLAB^®^ and used as nuisance regressors. The procedure removes unwanted sources of noise from the raw fMRI timecourse. At this point, the preprocessing pipelines were split. In total, three pipelines were investigated, a conventional SPM pipeline, a quasi-optimal SG pipeline suitable for shorter timecourses, an optimal SG pipeline suitable for longer timecourses. Data that underwent SPM detrending were either treated with a HRF based filter or a Gaussian smoothing kernel width = 2.48 sec (38). Data that were detrended with a SG (69/6) filter was subjected to a SG (15/8) low pass filter while data that was detrended with a SG (311/40) filter was subjected to a SG (3/1) low pass filter. The whole pipeline is depicted in Figure 6. The correct parameters of the SG filter used for detrending and high frequency noise removal were determined in a different working memory experiment described above.

**Figure 6:**
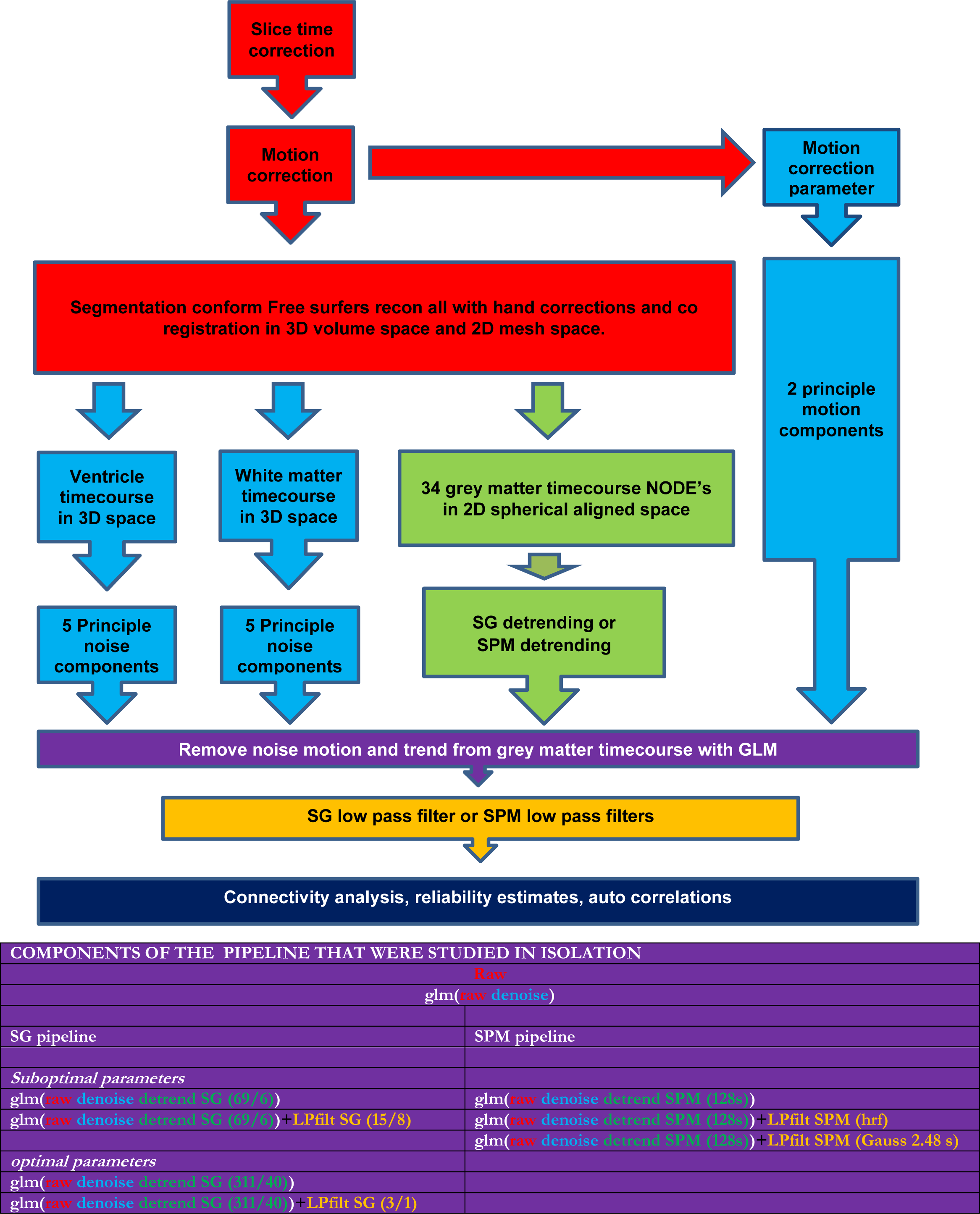
This figure gives the full preprocessing pipeline and a table that reports which components of the pipeline was studied in isolation. Red refers to the “raw data” part of the pipeline. Light blue refers to the denoising part of the pipeline. Green refers to the detrending part of the pipeline. Yellow refers to the low pass filter part of the pipeline. Purple refers to weighted noise removal within a GLM frame work. While dark blue refers to the analysis module. The colours of the letters in the table refer to the colours of the preprocessing depicted in the figure. Abbreviations, LPfilt=low pass filter; SG = Savitzky Golay. Results for the distinct preprocessing pipelines are given in Table 1-3

Our aim was to disclose whether SG filters performed better when compared to unfiltered data or classical SPM filters. We estimated all reliability and connectivity measures in a hierarchical ordered series of preprocessing steps. All relevant variables were simultaneously entered into the regression model except for the low pass filters that were applied after all other variables were regressed out. We report the preprocessing pipelines that were investigated in isolation in the table related to Figure 6. For reasons of readability we use the following nomenclature:

**Table 1:** This table reports observed and predicted grand mean autocorrelations of a test and retest run. The grand average was obtained by averaging the auto correlations of 34 nodes* 67 individuals. Observed autocorrelations were estimated from relevant timecourse sections for the various preprocessing methods mentioned in column 1 while predicted auto correlations were obtained from the constructed predictor timecourse that were based on denoised and SG (311/40) detrended data. RMSE = the root mean square error between predicted auto correlations and observed auto correlations.

|  | observed auto correlation |  |  |  | RMSE | observed auto correlation |  |  |  | RMSE |
| --- | --- | --- | --- | --- | --- | --- | --- | --- | --- | --- |
|  | test run |  |  |  |  | re test run |  |  |  |  |
|  | lag 1 | lag 2 | lag 3 | lag 4 |  | lag 1 | lag 2 | lag 3 | lag 4 |  |
| Raw | 0.66 | 0.48 | 0.36 | 0.24 | 0.22 | 0.66 | 0.48 | 0.37 | 0.24 | 0.22 |
| Denoised (5 wm+5 ventricle+2 motion components) | 0.56 | 0.34 | 0.21 | 0.09 | 0.24 | 0.58 | 0.37 | 0.24 | 0.12 | 0.24 |
| Denoised<br>Detrend SG (311/40) | 0.51 | 0.25 | 0.09 | -0.05 | 0.28 | 0.51 | 0.24 | 0.08 | -0.06 | 0.30 |
| Denoised<br>Detrend SG (69/6) | 0.51 | 0.25 | 0.10 | -0.04 | 0.28 | 0.52 | 0.25 | 0.09 | -0.05 | 0.30 |
| Denoised<br>Detrend SG (311/40)<br>Filtered SG (3/1) | 0.81 | 0.49 | 0.16 | -0.05 | 0.04 | 0.81 | 0.48 | 0.15 | -0.07 | 0.04 |
| Denoised<br>Detrend SG (69/6)<br>Filtered SG (15/8) | 0.79 | 0.40 | 0.08 | -0.08 | 0.08 | 0.79 | 0.40 | 0.07 | -0.10 | 0.10 |
| Denoised<br>Detrend SPM (128 s.) | 0.54 | 0.31 | 0.17 | 0.05 | 0.25 | 0.56 | 0.33 | 0.20 | 0.07 | 0.25 |
| Denoised<br>Detrend SPM (128 s.)<br>Filter SPM (HRF) | 0.92 | 0.73 | 0.49 | 0.24 | 0.38 | 0.92 | 0.74 | 0.51 | 0.27 | 0.38 |
| Denoised<br>Detrend SPM (128 s.)<br>Filter SPM Gaussian (2.48 s.) | 0.93 | 0.76 | 0.54 | 0.31 | 0.44 | 0.93 | 0.77 | 0.57 | 0.35 | 0.44 |
|  | predicted auto correlation |  |  |  |  | predicted auto correlation |  |  |  |  |
|  | re test run |  |  |  |  | test run |  |  |  |  |
|  | lag 1 | lag 2 | lag 3 | lag 4 |  | lag 1 | lag 2 | lag 3 | lag 4 |  |
| Denoised Detrended | 0.81 | 0.54 | 0.24 | -0.01 |  | 0.82 | 0.54 | 0.25 | 0.00 |  |

Raw data that underwent motion and slice time correction are referred to as “*raw*”.

“Raw” data that were denoised with motion, ventricle and white matter timecourses are referred to as “*denoised”.*

“Denoised” + detrended is referred to as “*SG or SPM detrended”.*

“SG or SPM Detrended” +low pass filtered (SG or SPM) is referred to as “*quasi-optimal or optimal SG filtered*” or *“Gaussian or HRF filtered”.*

### Conservative and liberal connectome

In this study every experimental trial was followed by a resting baseline condition. One might argue that the resting sections of the timecourse induce large signal changes that are not related to working memory in the narrow sense of the word. These large signal changes might increase the test retest reliability of a signal and therefore increase true connectivity. It is therefore reasonable to study how true connectivity among brain regions is changed when resting baseline sections are omitted from the timecourse. Thus connectomes were created with two approaches. First, we correlated the entire timecourse i.e. both the cognitive and resting baseline part of the timecourses. We refer to this as the liberal approach because it is likely to result in a relatively high true connectivity. Next, we correlated the section of the timecourses that were related to subtle changes in cognitive behaviour. We will refer to this as the conservative approach because it is likely to result in relatively low true connectivity estimates. We illustrate which section of the timecourse was used for the conservative approach on the basis of an event related average shown in Figure S1.

### Estimation of autocorrelations

We estimated the auto correlation structure of predicted and observed timecourses of every single node of every single individual in the spatial working memory experiment. The exact procedure that was used to obtain predicted and observed timecourses is described in the section “optimization experiment”. We assumed that the predicted timecourse should be based on 24 repetitions of an event related average that was obtained from a detrended fMRI signal because this preprocessing method is more or less standard. The autocorrelation structure of the predicted timecourse was compared with the autocorrelation structure of the observed timecourse sections. The test run served as predictor for the retest run and vice versa. In addition we estimated the lag1 autocorrelation of the high frequency noise as obtained after low pass filtering for all individual nodes per subject.

### Statistical policies

The purpose of this section is to describe how preprocessing strategies might affect the truly detectable connectivity between areas at the level of the individual which is closely related to the connectivity upper bound defined as:

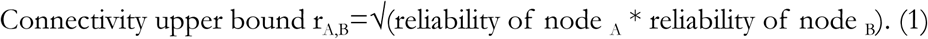

In the introduction it was shown that Formula 1 implies that reliability is estimated with Pearson correlations. In addition, it is common practice to estimate the connectivity between nodes with Pearson correlations and we will follow this tradition because connectivity can be anti-correlated. Treatment of timecourses within the GLM framework established yields inherently standardized timecourses. We *z*-transformed raw timecourses to ensure comparability of all preprocessing conditions. Discussions whether timecourse reliability should be obtained with ICC or Pearson correlations become superfluous (47) because ICC(1,1), ICC(2,1), ICC(3,1), and Pearson correlations performed over standardized timecourses yield practically identical point estimates. The very small differences observed are certainly not significant given the rather unreliable aspects of the measurements as such. It is thus safe to say that results reported generalize to all the mentioned correlation types. Fishers’ *𝒵* - transforms were performed whenever a correlation was subjected to some kind of statistical analysis such as averaging etc. We will use critical reliability thresholds as formulated by Cicchetti and count the number of individuals or brain connections that reach these critical thresholds (48). Cicchetti classified levels of ICC, in terms of practical or clinical significance, as follows: < 0.40 = Poor; 0.40 - 0.59 = Fair; 0.60 - 0.74 = Good; and > 0.75 = Excellent.

### Reliability and connectivity analysis of timecourses on the level of the path

#### Reliability

For every individual, test-retest reliability of 34 timecourses of interest was estimated per node leading to 34_reliabilities_*67_participants_ = 2278 correlations. The grand mean reliability correlation and the related confidence interval were estimated (Tables 3-4). Such correlations are difficult to interpret without reference. According to Cicchetti, correlations > 0.4 are at least fair; correlations > 0.6 are at least good; correlations > 0.75 are excellent. Using this logic, we estimated the following measures: First, we estimated the percentage of nodes that reached the critical reliability threshold per subject. Subsequently, we averaged this estimate over the whole sample and refer to this as "Mean percentage of nodes within-subjects”. Second, we calculated the average reliability per node leading to 34 correlations and estimated the percentage of nodes that reached the critical threshold and refer to this as "Percentage of mean nodes”. Third, we estimated the mean reliability of the 34 nodes per subject leading to 67 mean node correlations. We estimated the percentage of individuals whose mean node reliability reached the critical threshold and refer to this as “Percentage of participants with mean node reliability”. In addition we estimated the test retest reliability of the high frequency noise as obtained after low pass filtering for every node of every subject. Again grand mean reliability is reported.

**Table 2:** This table reports test retest reliability and connectivity statistics of working state data on the within subject level as a function of preprocessing method. Figure A reports the section of the timecourse that was used for the liberal correlation approach. Data were estimated from a connectome that consisted of 34 nodes obtained from 67 participants with a test retest design.

| Within subject within path reliability | Raw (slice time + Motion correction) | Denoised (5 wm 5 ventricle 2 motion) | Denoised Detrended SG (311/40) | Denoised Detrended SG (69/6) | Denoised DetrendedSG (311/40) FilteredSG (3/1) | Denoised DetrendedSG (69/6) FilteredSG (15/8) | Denoised Detrended SPM (128s.) | Denoised Detrended SPM (128s.) Filtered SPM (HRF) | Denoised Detrended SPM (128s.) Filtered SPM(2.48s.) |
| --- | --- | --- | --- | --- | --- | --- | --- | --- | --- |
| Grand mean connectivity estimate | 0.44 | 0.43 | 0.47 | 0.46 | 0.59 | 0.54 | 0.44 | 0.58 | 0.57 |
| Grand mean reliability estimate | 0.28 | 0.25 | 0.35 | 0.32 | 0.48 | 0.41 | 0.26 | 0.41 | 0.38 |
| Mean reliability upper bound | 0.36 | 0.33 | 0.42 | 0.40 | 0.54 | 0.48 | 0.34 | 0.48 | 0.45 |
| Mean reliability lower bound | 0.19 | 0.16 | 0.27 | 0.24 | 0.40 | 0.33 | 0.18 | 0.33 | 0.30 |
| Grand mean detectable connectivity maximum | 0.25 | 0.22 | 0.32 | 0.30 | 0.45 | 0.38 | 0.24 | 0.38 | 0.35 |
| Grand mean true connectivity | 0.22 | 0.22 | 0.31 | 0.29 | 0.43 | 0.37 | 0.23 | 0.37 | 0.34 |
| Absolute overestimation (r diff.) | 0.17 | 0.22 | 0.17 | 0.18 | 0.18 | 0.19 | 0.22 | 0.24 | 0.26 |
| Relative overestimation | 77 | 100 | 56 | 64 | 42 | 52 | 93 | 66 | 76 |
| % of corrupt paths | 15 | 8 | 3 | 4 | 4 | 4 | 5 | 5 | 6 |
| % of overestimated paths | 70 | 86 | 85 | 86 | 81 | 83 | 88 | 84 | 84 |
| Mean % of nodes within participants with (r>0.4) | 24 | 15 | 36 | 31 | 65 | 52 | 17 | 50 | 44 |
| Mean % of nodes within participants with (r>0.6) | 4 | 1 | 6 | 5 | 26 | 14 | 1 | 16 | 12 |
| Mean % of nodes within participants with (r>0.75) | 0 | 0 | 0 | 0 | 3 | 1 | 0 | 2 | 1 |
| % of mean nodes (r>0.4) | 3 | 6 | 21 | 18 | 68 | 50 | 9 | 50 | 41 |
| % of mean nodes (r>0.6) | 0 | 0 | 0 | 0 | 9 | 0 | 0 | 0 | 0 |
| % of mean nodes (r>0.75) | 0 | 0 | 0 | 0 | 0 | 0 | 0 | 0 | 0 |
| % of subj. with mean node reliability (r>0.4) | 7 | 1 | 15 | 7 | 70 | 52 | 1 | 46 | 40 |
| % of subj. with mean node reliability (r>0.6) | 0 | 0 | 0 | 0 | 1 | 0 | 0 | 1 | 0 |
| % of subj. with mean node reliability (r>0.75) | 0 | 0 | 0 | 0 | 0 | 0 | 0 | 0 | 0 |

**Table 3:** This table reports test retest reliability and connectivity statistics of working state data on the within subject level as a function of preprocessing method. Figure A reports the section of the timecourse that was used for the conservative correlation approach. Data were estimated from a connectome that consisted of 34 nodes obtained from 67 participants with a test retest design.

|  | Raw<br>(slice time<br>Motion<br>correction) | Denoised<br>(5 wm<br>5ventricle<br>2 motion) | Denoised<br>Detrended<br>SG<br>(311/40) | Denoised<br>Detrended<br>SG (69/6) | Denoised<br>DetrendedSG<br>(311/40)<br>FilteredSG<br>(3/1) | Denoised<br>DetrendedSG<br>(69/6)<br>FilteredSG<br>(15/8) | Denoised<br>Detrended<br>SPM<br>(128s.) | Denoised<br>Detrended<br>SPM<br>(128s.)<br>Filtered<br>SPM<br>(HRF) | Denoised<br>Detrended<br>SPM (128s.)<br>Filtered<br>SPM(2.48s.) |
| --- | --- | --- | --- | --- | --- | --- | --- | --- | --- |
| Grand mean connectivity estimate | 0.39 | 0.38 | 0.37 | 0.37 | 0.46 | 0.44 | 0.38 | 0.50 | 0.50 |
| Grand mean reliability estimate | 0.20 | 0.16 | 0.21 | 0.20 | 0.31 | 0.27 | 0.17 | 0.27 | 0.25 |
| Mean reliability upper bound | 0.31 | 0.27 | 0.32 | 0.31 | 0.41 | 0.38 | 0.28 | 0.37 | 0.36 |
| Mean reliability lower bound | 0.08 | 0.04 | 0.10 | 0.09 | 0.20 | 0.16 | 0.06 | 0.16 | 0.14 |
| Grand mean detectable connectivity maximum | 0.17 | 0.14 | 0.19 | 0.18 | 0.28 | 0.25 | 0.15 | 0.24 | 0.22 |
| Grand mean true connectivity | 0.14 | 0.13 | 0.18 | 0.17 | 0.26 | 0.23 | 0.15 | 0.23 | 0.21 |
| Absolute overestimation (r diff.) | 0.15 | 0.21 | 0.18 | 0.19 | 0.19 | 0.19 | 0.21 | 0.24 | 0.25 |
| Relative overestimation | 107 | 153 | 100 | 110 | 74 | 85 | 144 | 106 | 120 |
| % of corrupt paths | 26 | 15 | 8 | 8 | 7 | 8 | 11 | 14 | 15 |
| % of overestimated paths | 60 | 78 | 80 | 81 | 78 | 78 | 81 | 76 | 76 |
| Mean % of nodes within participants with (r>0.4) | 14 | 4 | 10 | 8 | 28 | 19 | 5 | 22 | 18 |
| Mean % of nodes within participants with (r>0.6) | 3 | 0 | 0 | 0 | 4 | 2 | 0 | 4 | 2 |
| Mean % of nodes within participants with (r>0.75) | 0 | 0 | 0 | 0 | 0 | 0 | 0 | 0 | 0 |
| % of mean nodes (r>0.4) | 0 | 0 | 0 | 0 | 12 | 0 | 0 | 6 | 0 |
| % of mean nodes (r>0.6) | 0 | 0 | 0 | 0 | 0 | 0 | 0 | 0 | 0 |
| % of mean nodes (r>0.75) | 0 | 0 | 0 | 0 | 0 | 0 | 0 | 0 | 0 |
| % of subj. with mean node reliability (r>0.4) | 1 | 0 | 0 | 0 | 15 | 6 | 0 | 9 | 6 |
| % of subj. with mean node reliability (r>0.6) | 0 | 0 | 0 | 0 | 0 | 0 | 0 | 0 | 0 |
| % of subj. with mean node reliability (r>0.75) | 0 | 0 | 0 | 0 | 0 | 0 | 0 | 0 | 0 |

#### Connectivity

The connectivities for test and retest run were computed for every single individual. In total 561connections are possible with 34 nodes leading to 561_connectivities_*67_participants_*2_measurements occasions_ = 75174 correlations. Subsequently, the grand mean connectivity coefficient was estimated from the 75174 correlations (Tables 3-4).

#### Detectable connectivity maximum

The detectable connectivity maximum was estimated for every single path for every subject on the basis of the 34 available reliability estimates according to Formula 1. Detectable connectivity maximum was considered a missing value when at least one reliability measure was negative. In total, 561_maxima_*67_participants =_ 37587 were estimated and subsequently averaged using the nanmean matlab command (see Tables 3-4). The latter command excludes nan values from the data

#### True connectivity

We averaged the connectivity correlations of the test and retest run per individual and refer to this as observed connectivity. We calculated true connectivity from the observed connectivity using following procedure.

> We set observed connectivity to nan when one or two nodes exhibit negative or zero test-retest reliability. We refer to this as corrupt connectivity.
>
> Otherwise if the observed connectivity is a positive correlation: In this case, we compared the observed and detectable upper bound correlations and took the smaller one.
>
> Else the observed connectivity is negative: In this case, we compared the absolute observed and absolute detectable upper bound correlations and took the smaller of the two and made the sign of the result negative.

We averaged true connectivity of every single path which represents the group average connectome consisting of 561 paths using nanmean command. The resulting mean true connectivity map was threshold at a fair r > 0.4 or good r > 0.6. We call this fair or good because the true connectivity map takes the underlying test-retest reliability of the path in question into account. Maps are shown in Figure 11-Figure 12. Finally, we estimated the grand mean true connectivity over paths and participants and report the result in Tables 2-3.

#### Absolute overestimation

The difference between the observed connectivity and the detectable maximum was estimated for every single path per subject when the observed connectivity was larger than the detectable maximum. Absolute overestimation was set at nan when et least one reliability measure was negative. The absolute overestimation was averaged over individuals and paths using the nanmean matlab command. In addition, we estimated the average percentage of overestimated paths per subject. Results are reported in tables 2-3.

#### Relative overestimation

We related the grand mean overestimation to the true grand mean detectable connectivity using the following formula:

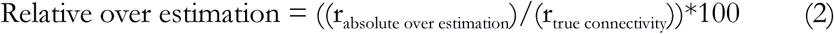

## RESULTS

### Behavioural data

Test-retest reliability of raw response time was excellent and in line with similar behavioural experiments executed during a stay in the scanner (49). Number Stroop task in spatial memory context ICC (2,1) = 0.75; Number Stroop task in verbal memory context ICC (2,1) = 0.83; spatial memory task ICC (2,1) = 0.77; verbal memory task ICC (2,1) = 0.78.

### Results of the Optimization experiment

#### Optimal detrending parameters

The optimization of detrending filter parameters was performed in the verbal working memory task. First, we wanted to know the extent to which correlation strength between predicted and observed timecourses is affected by SG parameters. Results of the exhaustive parameter search presented in Figure S2 reveals that data show irrational behaviour when polynomial orders larger than 42 are used. Hence, we zoomed in on the rational aspects of the data. The lines depicted in Figure 7 indicate an optimal family of SG parameters that is identified along a diagonal. Left of the diagonal a quasi-optimal SG parameter family is observed; right of the diagonal a disruptive SG parameter family is observed. Hence, the margin between optimal and poor SG parameter sets is very narrow. Differences in correlation strength of the parameter sets that belong to the quasi-optimal SG filter family are very small (Figure S3). Furthermore, SG parameters that belong to the quasi-optimal filter family exhibit linear scaling properties (figure S3). In our data, detrending was optimal with (311/40) SG filter. However, this filter calls for rather long experiments. We wanted to know if SG detrending is potentially effective in shorter experiment as well. Hence, for further analysis we also selected a quasi-optimal SG filter (69/6) that can be applied in experiments that are shorter in duration.

**Figure 7:**
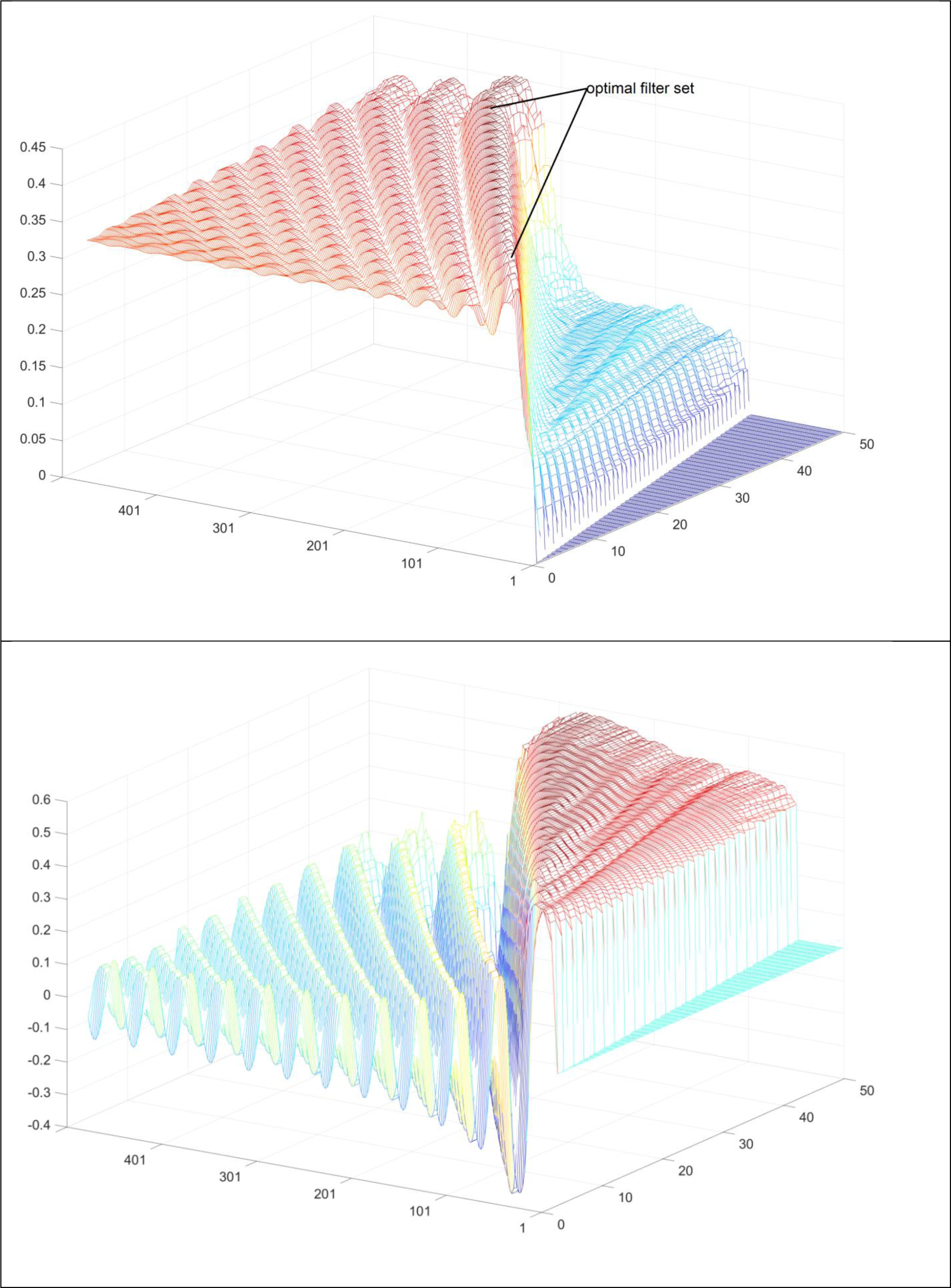
This figure depicts the average correlation between a predictor and observed timecourse for a given SG filter as obtained from 34nodes*67participants*2runs. The z axis represents the height of the correlation the left axis the size of the window (only uneven numbers) the right axis represents polynomial order. Mark that some filter parameters exhibit very small or even negative correlations Top: this graphs show results for a detrending filter that was developed in concert with denoising. Bottom: this graphs show the results for a low pas filter obtained after denoising and SG detrending (311/40).

#### Optimal low pass filter parameters

The optimization of low pass filter parameters was performed in a verbal working memory task. First, we wanted to determine how auto correlations of observed timecourse sections are affected by SG filters. Results of the exhaustive parameter search are depicted in Figure 8. It is clear that the curvature of lag 1 autocorrelations is far steeper when compared to curvatures of lag 2-4 autocorrelations.

**Figure 8:**
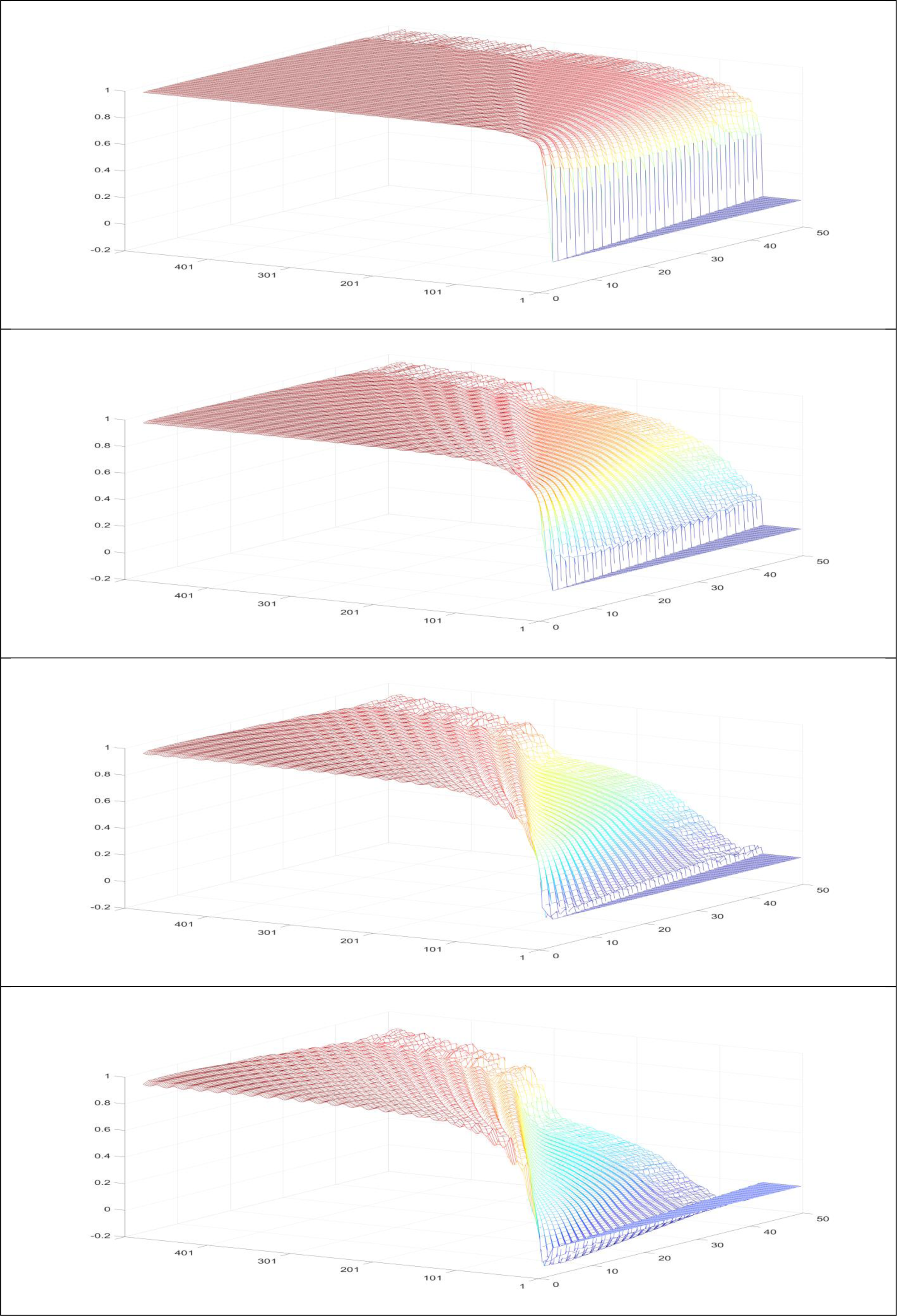
**This figure depicts the average autocorrelation of observed timecourses for a given SG filter as obtained from 34 nodes*67participants*2runs. From top to bottom the lag 1 to lag 4 auto correlations are represented. The z axis represents the height of the autocorrelation the left axis the size of the window (only uneven numbers) the right axis represents polynomial order. Mark wrong filters yield auto correlations that approach 1.**

In a next step, we estimated the lag 1-4 autocorrelations of the predicted timecourse. Ideally the auto correlation structure of the predicted timecourse should be similar to the autocorrelation structure of the observed timecourse. We estimated the root mean square error (RMSE) from the 4 autocorrelations of the predicted and observed timecourses for every single filter combination. An RMSE approaching zero reflects that observed timecourses that were treated with a particular SG parameter set exhibit similar autocorrelations as the predicted timecourse. Figure 9 depicts a family of SG filter parameters that approach the true auto correlational structure of the timecourses under study. This is indicated by the “valley” which approach RMSE values of 0. Subsequently, we wanted to know how the height of the correlation between predicted and observed timecourses is affected by SG parameters. The results depicted in Figure 7 complement results that are visualized for “detrending parameters”. In the case of low pass filters the margin between success and failure is even more dramatic and ranges from roughly 0.6 for effective filters down to -0.4 for ineffective filters. The mesh exhibits a triangular shaped plateau of maxima (Figure 9). We selected the correct family of low pass filters by masking the correlation mesh with the RMSE auto correlation mask. We only accepted RMSE < 0.1. The masking procedure led to a very well defined family of SG low pass parameters (Figure 9). Again, parameters that belong to the quasi-optimal filter family exhibit linear scaling behaviour (Figure S4). In simple words,

**Figure 9:**
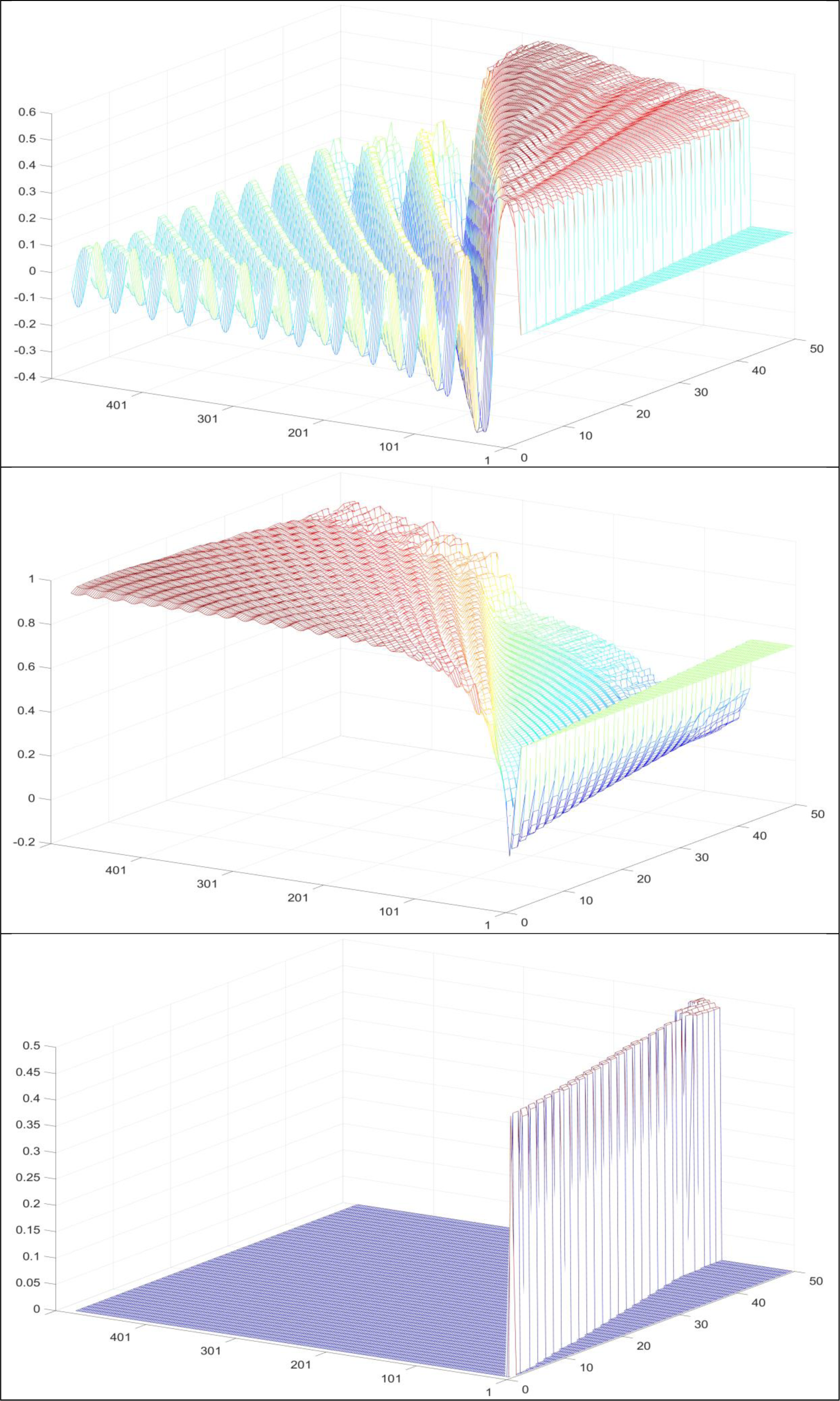
**This figure depicts the effect of SG high pass filter parameters on the correlation between predicted and observed timecourses; and the effect of SG parameters on RMSE obtained from predicted and observed autocorrelations. All data were obtained from 34 nodes *67participants*2runs. Predicted and observed timecourses underwent optimal SG detrending (311/40)** **Top: This figure is equivalent with Figure 7 (bottom panel). The z axis represents correlation height, the left axis the size of the window (only uneven numbers) the right axis represents polynomial order.** **Middle: The z axis represents RMSE the left axis the size of the window (only uneven numbers) the right axis represents polynomial order.** **Bottom: This figure shows what remains of the height of the correlation between predicted and observed data (top panel) when data are masked for RMSE<0.1 (middle panel).**

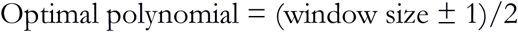

In our data, a low pass filter of SG (3/1) was optimal when it was combined with a detrending SG (311/40) filter. We also investigated a quasi-optimal low pass SG (15/8) filter that was combined with a quasi-optimal detrending SG (69/6) filter.

### Results of the validation experiment

#### Auto correlational structure of observed and predicted timecourses

In Table 1 we report the observed auto-correlation structures for the spatial working memory experiments as obtained with the various preprocessing methods and the autocorrelation structure of the predicted timecourse obtained from denoised data. The RMSE estimated from predicted and observed autocorrelations reveals that the closest match between predicted and observed auto-correlations is found when data underwent denoising, SG detrending, and SG low pass filtering. Results reported in Table 1 show that RMSE of the latter preprocessing types is very distinct from any other preprocessing type. This table also reveals that the autocorrelations of raw, denoised, and SG and SPM detrended timecourse sections are too low, while HRF and Gaussian filters yield autocorrelations that are much too high. One might criticize the use of denoised and detrended data as a gold standard because it is to a certain extent arbitrary in nature. Predicted timecourses that are based on other preprossessing methods show rather similar autocorrelation structures as long as SPM low pass filtering is not applied (Table S1). This is by no means surprising because the predicted timecourses were based on an event related average, which is in fact an effective way to denoise data. The latter is particularly true when event related averages have been obtained from jittered timecourses where systematic sources of noise are cancelled out. For instance, raw observed timecourses might suffer from high frequency noise, but the event related average of the raw timecourse does not suffer from high frequency noise because it is levelled out through the averaging procedure. Thus, while autocorrelations of observed timecourses react sensitively to differences in preprocessing, predicted timecourses do not because they are denoised through the averaging process and thus yield rather similar results irrespective of the preprocessing in use. The latter is however only true for timecourses that where not treated with aggressive low pass filters, as we will show in the next section. As an additional check we assessed the lag 1 autocorrelation of high frequency noise and its test retest reliability. Ideally the autocorrelation of noise timecourses should approach zero. Furthermore a test retest reliability of zeros suggests that the noise signal does not contain reliable cognitive information. Test retest reliability of high frequency noise as obtained by optimal and quasi optimal SG filters was on average 0.02 and 0.03 respectively. For HRF and Gaussian filters autocorrelation values of 0.06 and 0.08 were observed. Auto correlations of high frequency noise was -0.03 and -0.05 for the test and retest runs of the optimal SG filter while for the quasi optimal SG filter autocorrelations of -0.01 and -0.01 were observed. In analogy we observed values of -0.06 and -0.07 for the HRF filter and values of -0.10 and -0.11 for the Gaussian filter.

#### Properties of filtered timecourses

In this section we will show the effects of preprocessing procedure on power spectra of timecourses and the timecourses themselves. In a first step, we created the grand mean power spectra of the working memory network. Power spectra were obtained per node per subject. Power spectra of all individuals were subsequently averaged. This leads to a representative spectrum for the entire sample under study (Figure S5). As expected, all preprocessing methods show highest power at task frequency. Raw timecourses exhibit a peak in the lowest frequency range. In fact, the low frequency peak exhibits a density comparable with the task peak. The low frequency noise is effectively removed when timecourses are denoised with the timecourses of the white matter, ventricles, and head motion. The SPM high pass filter (128 secs. > 0.0078 Hz) shows almost no effect when it is combined with denoising methods. By contrast, SG detrending removes frequencies below task frequency. The lower frequencies that are removed through SG filtering are not slow trends that are in fact removed through the denoising operation. But they are possibly resting state signals that leak into the cognitive experiment. However, these unwanted zombie oscillations are certainly not related to the task of interest that is performed at higher speed.

We conclude that SG detrending methods effectively remove zombie oscillations - which are not related to our working memory task - from the timecourse while the latter is less the case for conventional SPM filter (128 seconds > 0.0078 Hz). The question arises if the observed effects of preprocessing on power spectra can be observed in the timecourses themselves. A closer look on a fraction of the timecourse of the working memory task reveals that denoising removes slow trends from the grey matter timecourse (see Figure S6). The benefit of a conventional SPM high pass filter is very little when it is combined with denoising methods (Figure S6). SG based high pass filtering has a slightly bigger effect on the timecourse (Figure 10 and S7). Figures 10 and S6-S7 might explain why SG filtering is more effective when compared to SPM filters. The conventional SPM filter only follows very rough trends, while SG filters capture oscillations that take place at much higher speed. Confirming the idea that SG based high pass filters act on a larger range of non-task related oscillations that may stem from resting state power spectra (Figure S5).

**Figure 10:**
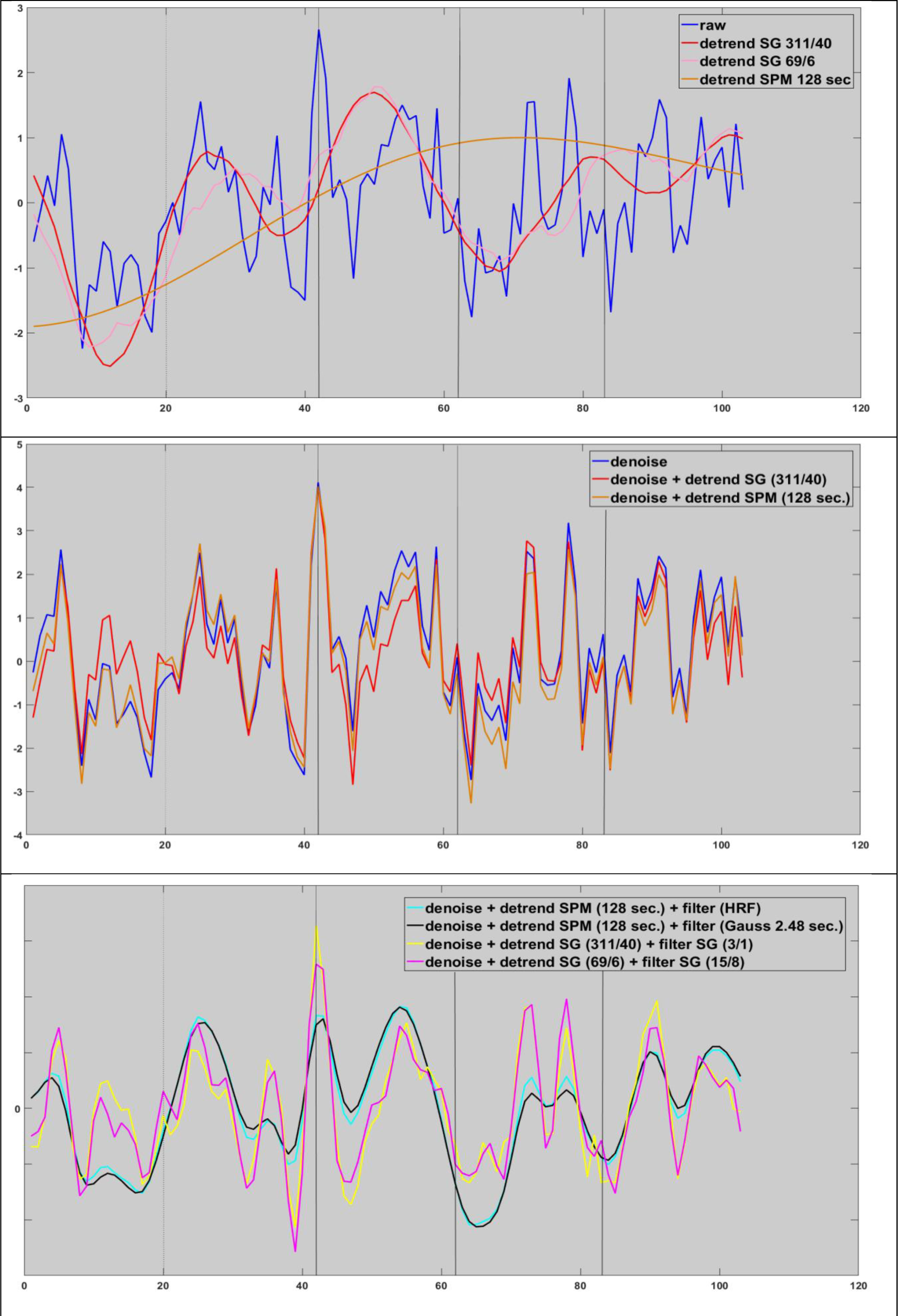
This figure depicts the effect of SPM and SG filters on an exemplary timecourse section. Top: conventional SPM filter detects very slow fluctuations while SG filters trace faster zombie oscillations. Middle: The effects of denoising and detrending executed within a GLM framework. Bottom effects of SPM and SG low pass filters Legends are given in the panel of interest.

**Figure 11:**
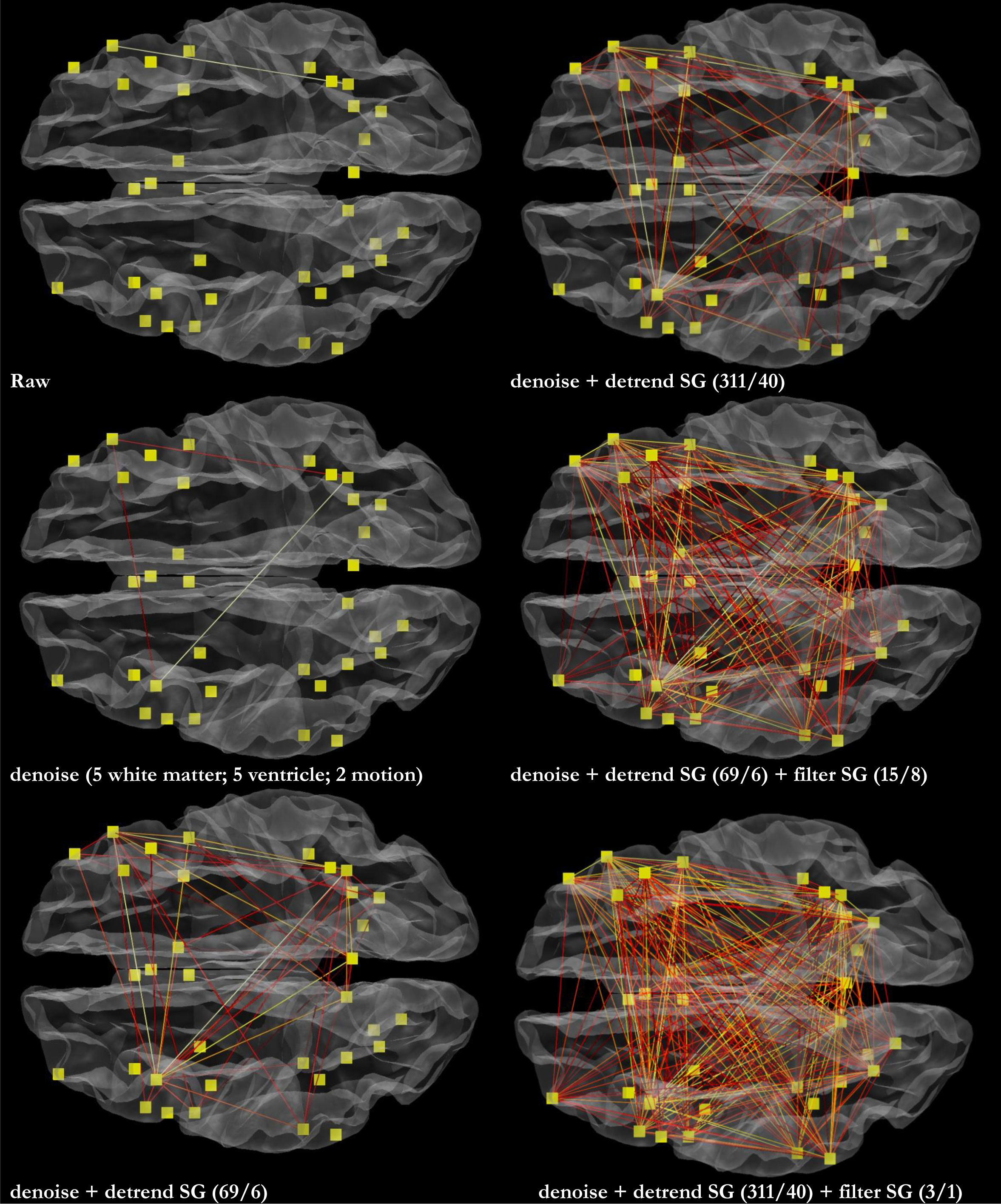
This figure reports how preprocessing affects true connectivity strength between 34 nodes of interest. Only connections with an average true connectivity of r>0.4 estimated from 69 individuals are shown. An average true connectivity of r>0.4 implies that average within subject timecourse reliability of the underlying nodes is at least fair r>0.4 Darker colours refer to lower correlations brighter colours to higher correlations.

**Figure 12:**
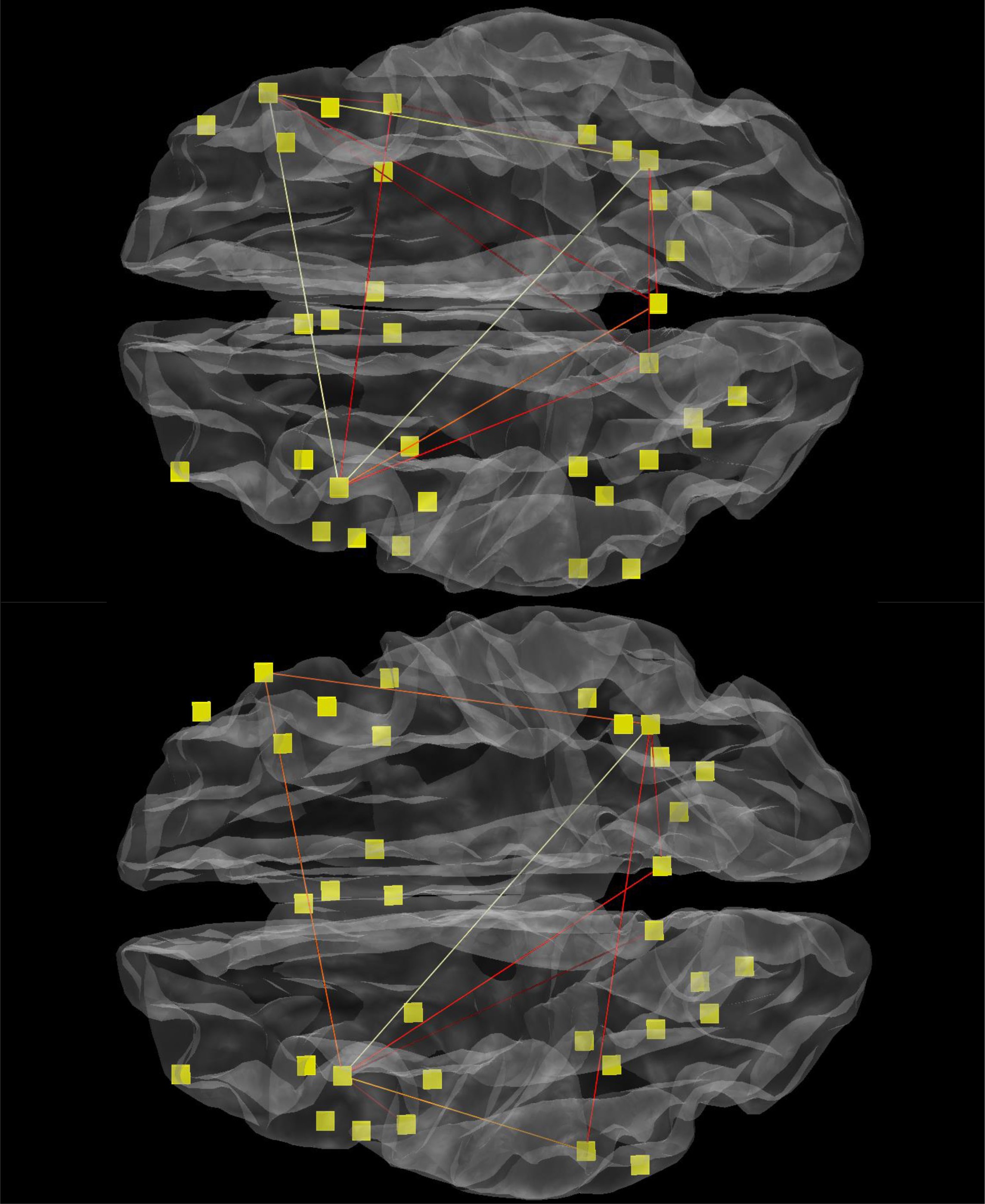
This figure shows the effects of correlation approach on the connectome of interest using an optimal SG pipeline. The liberal approach included cognitive action + rest while the conservative approach included cognitive action only. Top: Liberal correlation approach. Only paths with true connectivity > 0.595 are shown. Bottom: Conservative correlation approach. Only paths with true connectivity > 0.395 are shown. An average true connectivity of r>0.4 or r>0.6 implies that that average within subject timecourse reliability of the underlying nodes is at least fair r>0.4 or good>0.6.

In Figure 10 (see also S7) we compared quasi-optimal and optimal SG detrending on another timecourse fraction. It is obvious that the quasi-optimal SG (69/6) filter removes more fine grained trends when compared to the optimal SG (311/40) filter. It is possible that quasi-optimal detrending parameters result in a slightly overdetailed behaviour. The question arises whether SG low pass filters are also better as their SPM counterparts. The effects of SG and SPM low pass filters on detrended working memory data was gradual but substantial in nature as can be viewed from the power spectra depicted in Figure S5. The effects of HRF and Gaussian filters that were applied to the SPM detrended data were rather similar in nature and both reached their minimum at 0.15 hz. By contrast, SG high pass filters that were applied to SG detrended data reached their minimum around 0.25 hz. SG filters capture dynamic aspects of cognition that are obviously induced by our complex working memory task. A closer look on a fraction of the timecourse reveals that HRF and Gaussian filters destroy timecourse details while this is less the case for both SG low pass filters. (Figure 10 and S8). While SG high pass filters are sensitive to dynamics of signal as revealed by the shape of its timecourse, this is not at all the case for HRF and Gaussian filters. However, this figure also shows that there are limits to amount of detail that can be captured by SG filters. The effects of aggressive low pass filters is to a lesser extent observed in event related averages. Figure S1 illustrates that event related averages estimated from distinct preprocessing methods are rather similar in shape as long as low pass filtering is used in a conservative fashion. Aggressive low pass filters have a disruptive effect on the true shape of the event related average and abolish cognitively relevant information (Figure S1).

#### Reliability and true connectivity

The main purpose of this paper is to investigate if SG filters can be used to tackle the dubious charms of voodoo correlations on the within subject within path level. Hence, we will focus on true connectivity and the extent to which timecourse connectivity is overestimated given the underlying test retest reliability of timecourses.

Results reported in Table 2 and Table 3 indicates that true connectivity depends on preprocessing strategy and length of the segment of the timecourse investigated. We first present the results of the entire timecourse and then for the segments of the timecourses that were not related to rest. Table 2 reveals that lowest reliability and connectivity estimates were detected in denoised timecourses (mean connectivity = 0.43; mean reliability = 0.25) while optimal SG filtered data yielded the highest values (mean connectivity = 0.59; mean reliability = 0.48). This means that it is possible to double the height of the reliability estimate through advanced preprocessing. But although the latter is certainly true, we emphasize that confidence intervals of reliability estimates are remarkably alarming given the large number of time points (487) of the timecourses under study. Lower bounds of reliability estimate ranged from 0.16 for denoised data to 0.40 for optimally SG filtered data. Thus, even optimal preprocessing pipelines just manage to reach the fair reliability criterion when conservative lower bound point estimates of reliability are used

The mean percentage of nodes that could be detected with fair reliability (*r* > 0.4) within subjects was only 15% for denoised data but 65% for optimal SG filtered data. The percentage of nodes averaged over individuals that could be detected with fair reliability ranged from 3% for raw data to 68 % for optimally SG filtered data. Finally, the number of individuals whose average node system could be detected with fair reliability ranged from 1% for denoised data to 70% for SG filtered data. Results reported in Table 3 clearly indicate that within subject within path reliability reacts sensitive to the type of preprocessing in use, for true connectivity is closely linked to reliability and thus sensitive to differences in preprocessing method. Interestingly, the mean true connectivity is not affected by the denoising procedure. A true connectivity of r = 0.22 was detected in raw and denoised timecourses. Detrending within the regression context does not seem to affect true connectivity in a negative way. While SPM detrending does not improve true connectivity, SG detrending does (*r* = 0.29 and 0.31 for quasi-optimal and optimal SG filters). Not surprisingly, largest improvements in true connectivity were obtained when low pass filters were applied. Application of the HRF and Gaussian filters as available in the SPM packages leads to true average connectivities of *r* = 0.37 and *r* = 0.34. While application of quasi-optimal and optimal SG filters lead to true average connectivities of *r* = 0.37 and *r* = 0.43. So it is possible to double the true detectable connectivity through advanced preprocessing. The effects of preprocessing are clearly visible in the histograms depicted in Figure S9. The application of SG low pass filters leads to a substantial boost in connectivity and reliability and increases true detectable connectivity. However it should be mentioned that the percentage of overestimated paths at the within-subject within path level was severe. On average, 70-86% of the paths were overestimated depending on the preprocessing method in use. The relative overestimation was lower for preprocessing methods that involved some form of low pass filtering. The effects were rather substantial with relative overestimation percentages of 42-64% for SG filtered data as opposed to percentages of 66-100% for other preprocessing methods.

Next, we show how preprocessing might affect reliability and connectivity if one concentrates at the cognitive section of the timecourse in Table 3. In S1 we show sections of cognitive interest for all participants. As discussed in the method section, only these timecourse segments were used in the creation of conservative connectomes. Omitting the non-cognitive aspects of the timecourses has a devastating effect on all the measures reported in Table 3. Again, unfavourable reliability and connectivity estimates were detected in denoised timecourses (mean connectivity = 0.38; mean reliability = 0.16) while optimal SG filtered data yielded the highest values (mean connectivity = 0.46; mean reliability = 0.31). Again, lower bound confidence intervals were alarming and ranged from 0.04 for denoised data to 0.2 for optimally SG filtered data. This means that reliability of the conventionally denoised pipeline completely collapsed while reliability of optimal SG pipelines was poor (r<0.4)..

True connectivity of cognitive timecourse sections was much lower when compared to entire timecourses. True connectivity of denoised timecourses dropped from 0.22 to 0.13 while optimal SG filtered timecourses dropped from 0.43 to 0.26. Nonetheless, true connectivity of best preprocessing pipelines remained twice as high. Voodoo connectivity values ranged from 74 to 153 percent.

The increase of true connectivity with the increase of preprocessing steps with the liberal correlation approach is also visible for the connectome image as depicted in Figure 11. We decided to consider that average true connectivity of a specific path should at least reflect fair reliability (> 0.395). Moreover, given the high autocorrelations introduced by the SPM pipeline, we restricted the visualization of the connectome on non SPM preprocessing methods. The raw image shows only one fronto-parietal path. The denoised image shows a triangular constellation between the right parietal cortex and two frontal nodes. The detrended image shows the previously discussed fronto-parietal triangle with and additional network of fronto-parietal connections that mainly emerged from the right parietal cortex. In addition, frontal connection between the hemispheres emerged. The application of (quasi) optimal SG filtering leads to an enormous increase in overall connectivity including left parietal and frontal polar systems. The optimal SG pipeline contained a small connectome that exhibited good (r>0.6) true connectivity (Figure 12). According to the conservative correlation approach, only the connectome of the optimal SG filter pipeline reached the critical average true connectivity threshold of 0.395. The connectome depicted in Figure 12 shows that the middle frontal gyrus (MFG) is the most important hub of the connectome. The MFG is roughly equivalent to the dorso lateral prefrontal cortex. This area is believed to be the core neural correlate of working memory processes (46). Working memory researchers usually claim that an area is relevant for working memory when it exhibits sustained brain activity during the course of the experiment. The graph depicted in Figure S1 that was based on the optimal SG filtered pipeline suggests that the latter is the case for most individuals under study.

### Simultaneous filter and nuisance regression

Previous research on resting state data suggested that noise is reintroduced into the timecourses of interest when filters are applied after denoising (50). Hence, our pipeline can be criticized because we indeed optimized and validated low pass filters after denoising and detrending data within a GLM framework. However it should be said that bandpass filters as applied in resting state research are not suitable for cognitive research because they may corrupt cognitive signal of interest. We investigated what happens with the signal if low and high pass filters as well as nuisance timecourses are simultaneously introduced into the GLM. We observed that the effects were strikingly different for the distinct filter types under study. RMSE values reported in Table S2 suggest that autocorrelations of the SPM pipeline dropped substantially when low pass filters were integrated into the GLM. The better temporal resolution led to a slight loss of true detectable connectivity (Table S3). Altogether, SPM filters when applied at all should be integrated into the GLM and not run after preprocessing. But autocorrelations and relative over estimation estimates of SPM low pass filtered data remain very problematic independent of the pipeline in use. Hence, we did not consider these pipelines in further analysis. Regarding SG filters, results where more puzzling. We again optimized a filter in the verbal working memory task using brute force methods. But as discussed this time all parameters were entered into the GLM at once. This leads to a SG low pass filter with a window size of 105 and a polynomial order of 35. Validation of the filter in the spatial working memory data led to a substantial boost in true detectable connectivity. Non intergraded SG filters exhibit an average true connectivity of 0.43 while integrated filters exhibit a true connectivity of 0.48 (Table S3). In addition, no less than 94% of individuals exhibit a mean node reliability of 0.4 when integrated low pass filters were used whereas this number was only 70% for non-integrated low pass filters. Superficially integrated SG filters seem to do a better job. However the price for this is a substantial increase in temporal autocorrelation (Table S2). In this context RMSE of non-integrated filters was roughly 0.05 whereas integrated approaches exhibited an RMSE of 0.15. The potentially destructive effects of the integrated approach on the timecourse of interest can be seen in the power spectra shown in Figure S10.

## DISCUSSION

### SG filters improve true connectivity while conserving realistic auto correlations

A previous task driven fMRI study reported that within subject reliability of timecourses may not exceed 0.25 (22). This suggests that true connectivity among region A and B cannot exceed 0.25. We have investigated if the hypothesized maximum value of 0.25 can be observed in experimental data. We used a similar pipeline as our colleagues .i.e. removed noise through regression analysis and high pass filters (128 seconds). Grand mean true connectivity of a working memory related connectome consisting of 34 nodes measured in 67 participants was very poor (r = 0.23). This is somewhat puzzling because observed connectivity among the very same regions was 0.44 meaning that roughly half of the observed connectivity variance is voodoo. We tried to improve the reliability of the timecourses with classic SPM low pass filters. The latter included a Gaussian filter (2.48 seconds) and a HRF filter. Application of these filters led to a substantial boost in connectivity. But these classic filters also boosted autocorrelations substantially. Observed lag 4 autocorrelations were roughly 0.3 meaning that classic SPM low pass filters may obscure temporal information of timecourses. We conclude that low pass filters have been omitted from the classic SPM package for good reasons. Results of SG low pass filters were more favourable. We observed a fair true connectivity of 0.43 and a connectivity of 0.59 in the optimal SG based preprocessing pipeline. Generally these findings suggest that roughly one third of the observed connectivity variance is voodoo. The percentage of nodes that were measured with at least fair (r > 0.4) or at least good reliability (r > 0.6) at the within subject level was on average 65% and 26% respectively. Furthermore, application of the SG based pipeline results in grey matter timecourses that approach the auto correlational structure of the idealized bold response. This observation was complemented by an analysis that focused on the auto correlation of high frequency noise. As expected autocorrelation of SG filtered noise signals was very low. Remarkably, autocorrelations of noise time courses was lower for quasi optimal (r=0.01) than for optimal (r=0.04) SG filters. But as discussed in the introduction the assumption that fMRI noise approaches idealized noise must not hold true. The temporal resolution of SG filtered signals is relatively high because bold fluctuations reflect dynamic changes in cognitive behaviour. This may be relevant fc-MRI applications that investigate rapid changes in connectivity patterns. Dynamic connectivity studies might be feasible because novel imaging methods may trace neural effects at a speed of 0.75 Hz (51). In this study timecourses of a predefined patch of interest were averaged. But colleagues have shown that it is better to average timecourses of neighbouring regions that share common temporal variations (17). Maybe it is best to combine this averaging approach with an SG filter.

### SG filters detect zombie oscillations

SG filters that are designed to detrend MRI signals improve within subject timecourse reliability while this is not the case with the classic SPM high pass filter (128 seconds). Our analyses suggest that SG filters detect oscillations not related to task, which are substantially faster than oscillations removed through classic SPM high pass filters. We can exclude the possibility that zombies are related to scanner drift, physiological noise as captured by our regressors or head motions. However, exclusion is in this case not a sufficient method to achieve a clear definition of what zombies are. We can only speculate about the biological meaning of zombie oscillations. Zombie oscillations may contain physiological noise that was not removed by the standard denoising method. Furthermore, Figure S5 shows that SG filters suppress oscillations that originate from the “resting state frequency band”. Hence, one might speculate that resting state signals contaminate working state signals(45). As determined in the analyses, zombie oscillations are too fast for ordinary signal trends. Hence, zombie oscillations may reflect brain activity that is not under the voluntary control of the individual. These unreliable forms of brain activity may interfere with working state signals. We called these oscillations “zombie oscillations” in analogy to the undead neural process described in a paper entitled “The zombie within” by the late Nobel laureate Crick (52). But whether zombie oscillations in the latter sense of the word exist is subject of future research.

### Reliable detection of subtle bold changes require refined pipelines

In the present study, we removed resting baseline sections from the fMRI timecourse. This yielded an analysis that focused on subtle task induced oscillations in the 0.05-0.2 Hz range. Average true connectivity of denoised timecourses dropped from 0.22 to 0.13 while true connectivity of optimal SG filtered timecourses dropped from 0.43 to 0.26. None of the pipelines were able to detect the overall connectome with a grand mean timecourse reliability of r > 0.4. However figure 12 shows that timecourses that were treated with optimal SG filters yielded an interpretable connectome that exhibited sufficient within subject timecourse reliability (r>0.4) at the node level. We conclude that experiments that focus on subtle changes in brain activity might require refined pipelines. Currently, we lack hard prove that conservative connectivity estimates that exclude the resting baseline from the timecourse are indeed better when compared to liberal connectivity estimates that include the resting baseline. In fact removing the baseline form the timecourse may introduce “cutting artefacts”. It has been suggested that cognitive aspect of the BOLD response should be removed from the signal trough covaraince analysis or low pass filters. Superficially this seems to be a valid method because it induces stationarity without artefacts. In our opinion it is rather speculative to assume that the remaining signal indeed reflects cognitive behaviour. Moreover our pipeline effectively removes most signals that are not related to the cognitive signal of interest as can be viewed from power spectra presented in Figure S5. Thus removing the cognitive aspect from the time course may result in a rather empty timecourse.

### Low pass filter should be used after denoising

Previous resting state studies claim that overestimation of connectivity is reduced when Fourier filtered signals and nuisance factors are simultaneously regressed out within a GLM framework (50). In this task driven connectivity study that focused at BOLD oscillations in higher frequency bands the opposite was true. SG low pass filters integrated in a GLM frame work yield slightly higher absolute over estimation values when compared to SG filters that were run over timecourses after preprocessing. For this reason, we do not recommend to integrate SG filters in a GLM framework although they may boost true connectivity substantially. We conclude that it is not possible to generalize the previous observations that have been obtained from Fourier based filters to SG filters (50). We suggest that aggressive bandpass filters - that are frequently used in resting state studies - should be regarded with scepticism because they may remove cognitive and emotional information from the timecourse.

### Limits of pipeline optimization

Tables 2 and 3 reveal that improvements made by SG filters are substantial indeed. This is not only true for optimal SG pipelines but also true for quasi optimal SG pipelines that can be run over shorter timecourses. But we do not know if the limits of true connectivity maximization have been reached by our SG pipeline or if further improvements are possible. One might argue that within subject within region reliability is a very conservative measure because humans inherently exhibit response variability that is reflected in the timecourse. Consequently, within region timecourse reliability is not only limited by the scanner or preprocessing method in use but also by brain activity itself. Currently it is difficult to distinguish response variability from noise. Thus even though psychologically induced timecourse variability might exist, it remains a speculative and hard to measure phenomena in the context of fMRI.

### Are fc-MRI pipelines alternative map machines?

In the past, function related brain areas that were detected with classic BOLD experiments were discussed within the context of neuropsychological lesion studies. In addition, multi method experiments suggested that the BOLD response might have a true neural basis (53). This gave classic bold imaging some credibility. However, a comparative investigation of over 6000 fMRI pipelines shows that classic BOLD imaging is not unproblematic (54). Researchers are confronted with a large number of alternative maps that may or may not reflect biological reality (54). The same argument holds of course true for connectivity studies. In the present study, more than 100000 fMRI pipelines were tested to find an optimal SG filter combination. The number of alternative pre-processing methods may easily run into the billions when diverse filter methods are combined with variable numbers of noise variables that may have been obtained through various methods. Here, we focus on 4 pipelines that survived the filter optimization procedure. We concentrate on the number of paths that exhibited a fair average true connectivity > 0.4 in face of acceptable autocorrelations. The “raw” pipeline yielded only one path, a pipeline that combined noise regression with an SPM high pass filter yielded 4 paths. A pipeline that combined noise regression with SG detrending yielded 74 paths. Adding an SG low pass filter to the latter pipeline resulted in a true connectome of 352 paths. The differences between the connectomes depicted in figure 11 are substantial. The question is which connectome reflects biological reality? Unfortunately, we don’t know because we only have statistical arguments pro or contra a pipeline. Hence causal external validation of fc-MRI pipelines with methods other than fMRI is required. Currently, we do not have connectivity studies that compare the effects of distinct MRI signal processing pipelines in combination with invasive direct brain stimulation and direct brain recoding techniques. This is remarkable because these kind of experiments are feasible (55) (56). However, multi method experiments are challenging because distinct imaging modalities may leave their traces in the signals of interest. Alternatively, one might try to improve fMRI pipelines on the basis of computer simulations. The advantage of the simulation method is that ground truth can be known. But computer simulation methods are based on the idea that realistic noise simulation is feasible. However one might question if white, pink or other forms of synthetic noise are true approximations of real world noise. In fact the autocorrelation of the noise timecourse was not zero for the optimal SG pipeline. Hence fMRI high frequency noise is possibly not equivalent with white noise. Moreover zombie oscillations are possibly difficult to simulate since they might contain psychological variability of unknown origin. Finally, it is likely that some sources of fMRI noise are still waiting to be discovered (45).

Problems around the validity of fc-MRI draw a sombre light on its clinical relevance. To the best of our knowledge no fc-MRI pipeline has been approved by the US Food and Drug Administration for clinical use (https://www.fda.gov/medicaldevices/). This remarkable since the FDA approved several classic BOLD imaging software packages as diagnostic instruments for invasive medical purposes. Thus the use of fc-MRI in diagnostic settings remains controversial. In contrast to structural MRI, fc-MRI never reached the status of a standard clinical routine in neurosurgery. Currently, most neurosurgeons still rely on direct brain stimulation methods that provide a causal link between a specific brain area and a cognitive function.

### Limitations of this study

This study is possibly the first fc-MRI study that investigates the effects of signal processing on true connectivity. We are left with the platitude that one study is no study. In the present study, SG filters were developed for slow event related design. It is very unlikely that our pipeline generalizes to other fMRI experiments. However, with a few further studies it may be possible to derive a function connecting the timing parameters of fMRI experiments, noise sources and the adequate SG filters necessary to deal with them. We speculate that optimal fMRI pipelines parameters may differ from experiment to experiment, brain region to brain region and individual to individual. However we were unable to show this. Furthermore we did not investigate if different repetition times might affect SG filter parameter optimization. In addition we did not study if behaviour of classic SPM filters can be improved when parameters are changed for modern purposes. It was beyond the reach of this study to compare SG pipelines with principal- or independent component analysis, wavelet analysis or autoregressive integrated moving average methods. In principle these methods although complicated may outperform our SG based method. We took the participants out of the scanner and repeated the measurements after some time. Thus, the head position and physiological condition of the participants were changed. But it is possible that reliability of our study is overestimated because test and retest sessions were obtained at the same day (18). The power spectra presented in Figure S5 suggest that SG filters remove noise in a very effective way. However, we did not obtain sources of physiological/psychological noise such as heart rate, respiration rate, electrodermal activity, body temperature, or eye motion. We think that it is likely that these unwanted sources of noise were removed by our pipeline but we have no proof of this. Finally one could argue that SG filters may also improve reliability of sophisticated methods such as granger causality, structural equation modelling or dynamic causal modelling. But this was again not within the reach of this study. One could use reliability estimates to correct connectivity estimates for measurement error (24). In fact we have used this kind of methods in a genetic context in a previous study (30). However we feel that the time is ripe to show the unadorned reality of fMRI.

## CONCLUSION

So far, more than 40.000 fMRI papers have been published but very few papers report test retest reliability of timecourses at the within subject within region level (11, 22). This is remarkable since timecourse analysis is at the core of every fMRI method. We observed that Savitzky Golay (SG) filters simultaneously maximize true connectivity among brain regions and minimize spurious auto correlations of timecourses. However the use of SG filters is not without risks since ill designed SG filters have an extraordinary destructive effect on the reliability of the signal. We used Cicchetti judgment values to interpret our results. In this context, poor timecourse reliability estimates (mean reliability under bound r < 0.4) were classified as insufficient for scientific purposes. In the present study, only SG based pipelines were on average sufficiently reliable for scientific purposes (mean reliability under bound r > 0.4). The road to timecourse reliability of clinical quality (r > 0.75) might be long. We observed that roughly one third of the observed connectivity variance remains voodoo despite pipeline optimization. Unfortunately, we do not know if the favourable results obtained from SG pipelines are generally valid because we lack external validation with a method other than fMRI. It is quite possible that other pipelines with poor statistical properties are better suited to display neural reality. Thus, SG pipelines may add to the confusing amount of existing pipelines. Timecourse reliability is possibly a very conservative reliability measure given the poor signal to noise ratios of fMRI and the inherent response variability of participants. In addition time course reliability of resting state data cannot be estimated. In this context it may be legitimate to measure reliability of fMRI on a group level that result in substantially higher reliability. Whether these relaxed reliability estimates lead to single subject connectomes suitable for invasive medical purposes remains to be seen.

Our study shows that it is fairly easy to manipulate fc-MRI data but fairly difficult to find the truth in fc-MRI data. Fc-MRI may suffer from a validity crisis when causal external validation of fc-MRI pipelines is executed at current rate.

## Ethics

All individuals provided informed consent. The research was executed according to the guidelines of the Declaration of Helsinki. The study was approved by the Ethics committee of the University of Graz under GZ 39/31/63 ex 2011/12.

## Contributions

J.W.K.: Conceptualization of SG preprocessing pipeline and voodoo connectivity analysis, programmed computer code, wrote rough draft of manuscript.

A.S.: Improved design of SG pipeline substantially, expert opinion on computer code, corrected rough draft.

V.K.: Projected meta-analysis coordinates into FreeSurfer Mesh space

G.W.: Approved and corrected rough draft.

